# Airway Basal Cells show a dedifferentiated KRT17^high^Phenotype and promote Fibrosis in Idiopathic Pulmonary Fibrosis

**DOI:** 10.1101/2020.09.04.283408

**Authors:** Benedikt Jaeger, Jonas Christian Schupp, Linda Plappert, Oliver Terwolbeck, Gian Kayser, Peggy Engelhard, Taylor Sterling Adams, Robert Zweigerdt, Henning Kempf, Stefan Lienenklaus, Wiebke Garrels, Irina Nazarenko, Danny Jonigk, Malgorzata Wygrecka, Denise Klatt, Axel Schambach, Naftali Kaminski, Antje Prasse

## Abstract

Idiopathic pulmonary fibrosis (IPF) is a fatal disease with limited treatment options. In this study we focus on the profibrotic properties of airway basal cells (ABC) obtained from patients with IPF (IPF-ABC). Single cell RNA sequencing of bronchial brushes revealed extensive reprogramming of IPF-ABC towards a KRT17^high^ PTEN^low^ dedifferentiated cell type. In the 3D organoid model, compared to ABC obtained from healthy volunteers, IPF-ABC give rise to more bronchospheres, *de novo* bronchial structures resembling lung developmental processes, induce fibroblast proliferation and extracellular matrix deposition in co-culture. Intratracheal application of IPF-ABC into minimally injured lungs of Rag2^-/-^ or NRG mice causes severe fibrosis, remodeling of the alveolar compartment, and formation of honeycomb cyst-like structures. Connectivity MAP analysis of scRNA seq of bronchial brushings suggested that gene expression changes in IPF-ABC can be reversed by SRC inhibition. After demonstrating enhanced SRC expression and activity in these cells, and in IPF lungs, we tested the effects of saracatinib, a potent SRC inhibitor previously studied in humans. We demonstrated that saracatinib modified *in-vitro* and *in-vivo* the profibrotic changes observed in our 3D culture system and novel mouse xenograft model.

## INTRODUCTION

Idiopathic pulmonary fibrosis (IPF) is a progressive disease with a lethal prognosis despite the introduction of new antifibrotic treatments ^1^. The histological pattern of IPF, usual interstitial pneumonia (UIP), is characterized by patchy and peripheral remodeling of the alveolar compartment ^2, 3^ with replacement of the normal alveolar architecture by fibroblastic foci, honeycomb cysts, and distorted airways. Recent data indicate that IPF exhibits features of small airway disease ^4^. Bronchiolization, the replacement of the resident alveolar epithelial cells by ABCs within remodeled regions in the IPF lung ^5-10^, leads to a dramatic shift in the epithelial cell repertoire of the alveolar compartment. The ABC is the progenitor cell of the airway epithelium and can give rise to any type of airway epithelial cell such as secretory, goblet or ciliated cells ^11^. ABCs play an important role in lung development, start the branching and tubing morphogenesis by building up the airway tree ^12^, and have been implicated in the pathogenesis of chronic obstructive pulmonary disease and lung cancer ^13-15^. We have recently confirmed the abundance of ABCs in the lung parenchyma of patients with IPF and discovered that the presence of an ABC signature in the transcriptome of bronchoalveolar lavage (BAL) cell pellet of patients was indicative of enhanced disease progression and mortality ^16^. Recent scRNAseq data of IPF tissues confirmed increase in ABCs of IPF lung tissues and described a unique aberrant basaloid KRT17^+^ cell population, which lacks the characteristic basal cell marker KRT5 ^8, 17, 18^. However, none of these studies tested functional properties of human IPF-ABC. Up to date, it has not been clear whether ABCs contribute to the fibrotic process or simply represent a bystander phenomenon.

In this study we sought to determine the transcriptional changes of IPF-ABCs and their potential profibrotic properties using several translational models. Single cell RNA sequencing of bronchial brushings demonstrated extensive reprogramming of IPF-ABCs towards a dedifferentiated KRT17^high^ PTEN^low^ cell type with high expression of various transcription factors associated with stemness. Using the organoid model, we determined that IPF-ABCs obtained from individuals with IPF (IPF-ABCs) are characterized by significantly increased formation of bronchospheres, *de novo* bronchial structures resembling lung developmental processes, and by having profibrotic properties on lung fibroblasts. Using a novel *in-vivo* xenograft model, we discovered that ABCs from individuals with IPF have the capacity to augment fibrosis, remodeling and bronchialization in the mouse lung. Connectivity map analyses revealed enrichment of SRC signaling in IPF-ABCs. Saracatinib, a known inhibitor of the SRC kinase pathway, blocked bronchosphere generation *in-vitro* and attenuated bronchialization and fibrosis *in-vivo*.

## RESULTS

### Single cell sequencing of bronchial brushes identifies reprogramming of IPF-ABCs in contrast to ABCs derived from disease control (NU-ABC)

Single cell RNA sequencing (scRNASeq) was performed on cells obtained by bronchial brushing from nine IPF patients and six nonUIP ILD controls (Table S1). 14,873 epithelial single cell transcriptomes were profiled. Based on expression profiles of known marker genes we identified four epithelial cell populations in the brushed cells (Figure 1A-C and Figure S1): ciliated cells (FOXJ1, HYDIN, 41.5% of all epithelial cells), secretory cells (SCGB1A1, SCGB3A1, 10.5%), ABCs (TP63, KRT5, 47.1% of cells) and ionocytes (STAP1, PDE1C, 0.8% of cells) and focused our analysis on the basal cells.

**Figure 1.**
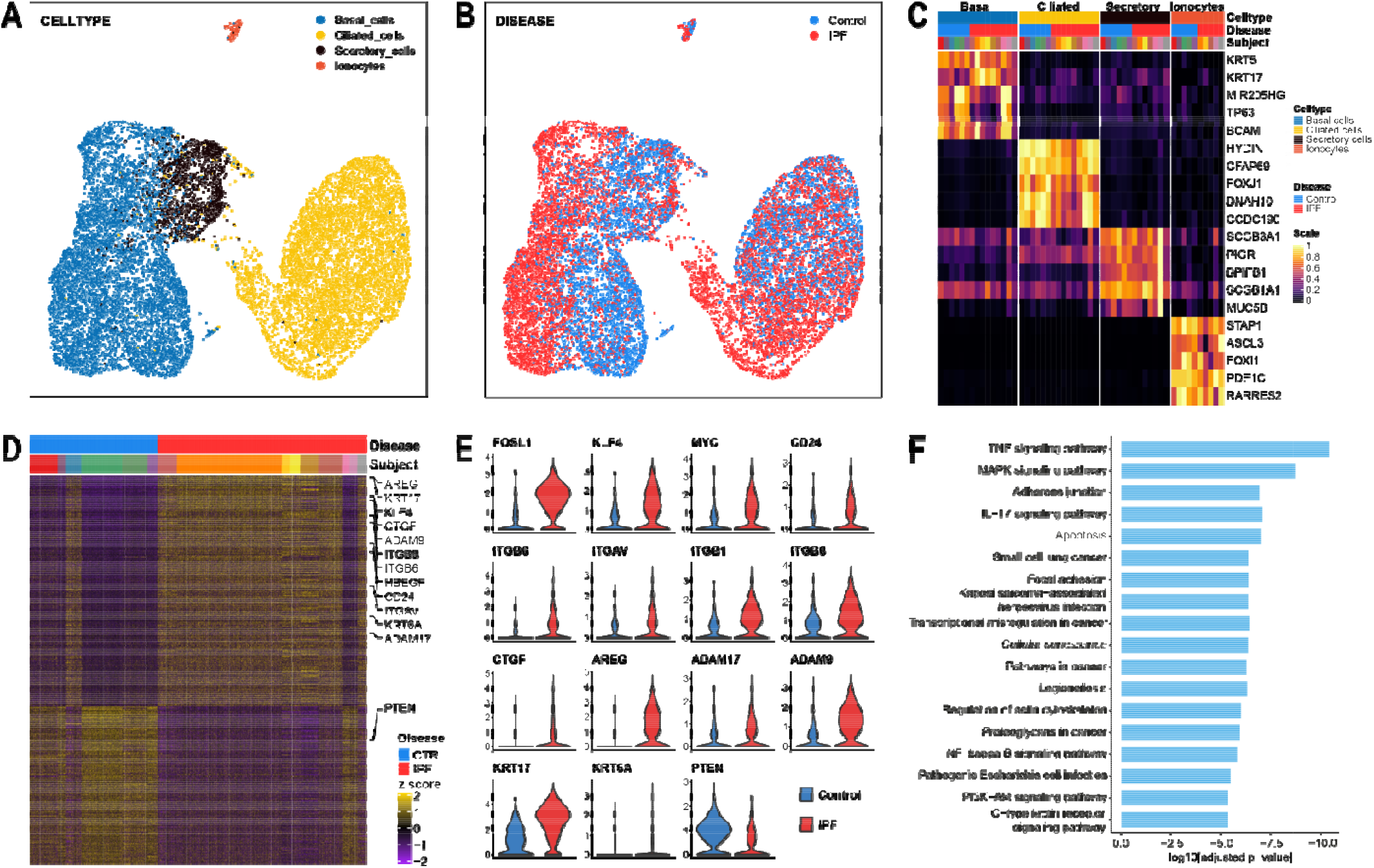
Single cell RNAseq and Clue analyses of bronchial epithelial cells. Single cell RNA sequencing was performed on Brush cells of IPF patients (n=9) and nonUIP ILD controls (n=6). Uniform Manifold Approximation and Projection (UMAP) of 14,873 single cell transcriptomes visualizes the four major discrete epithelial cell types (detected UMAPs colored by **(A)** cell types, **(B)** disease state). **(C)** Heatmap of unity normalized canonical epithelial m rker gene expression, averaged per subject, grouped by cell type, as shown in (A). **(D)** Heatmap of differentially expressed genes in ABCs of IPF patients vs nonUIP ILD disease controls showing a distinct deviation of gene expression (each column is a ABC, each row a gene as z-scores), **(E)** Violin plots of a subset of differentially expressed genes (DEGs) split by disease state. **(F)** Bar plot of log10-transformed adjusted p-values from the pathway analysis using the human KEGG 2019 database.

Gene expression of IPF-ABC was substantially different from NU-ABC (1,099 genes at FDR<0.05, Figure 1D). The IPF basal cell subpopulation was characterized by an increased expression of stem cell markers and stemness increasing signal transduction factors such as FOSL1, KLF4, MYC, CD24 which was accompanied by a loss of PTEN (Figure 1D and E). Expression of KRT17, KRT6A, stratifin (SFN), CTGF and genes for integrin subunits such as ITGB6, ITGAV, ITGB1 and ITGB8 were highly upregulated (Figure 1D and E). Moreover, an increased expression of the EGF family members AREG and HBEGF and the shedding enzymes for amphiregulin, ADAM17 ^19^, and for EGFR, ADAM9 ^20^, was observed (Figure 1D and E). Pathway analyses showed multiple pathways upregulated related to cancer, mechanotransduction, ECM sensing, cellular senescence, EMT, TNF-α and IL-17 signaling (Figure 1F). Of note, we observed a considerable overlap to the recently described pathways which are upregulated in KRT17^+^ aberrant basaloid cells ^17, 21^.

### ABCs from IPF patients generate significantly more bronchospheres than ABCs from healthy volunteers or nonUIP interstitial lung disease (ILD) patients in a 3D organoid model

While cell proliferation of IPF-ABCs compared to those obtained from healthy volunteers (HV-ABC) or individuals with fibrotic nonUIP ILD (NU-ABC) did not differ in 2D cell culture, the bronchosphere formative capacity in a 3D organoid model was very different (Figure 2A-D). After 21 days, all three cell types formed organoids in 3D culture, however IPF-ABC formed organoids that were multilayered and frequently contain hollow and tube-like structures, visually resembling bronchospheres (Figure 2B and C). The average number of spheres per well generated by IPF-ABCs was significantly higher compared to HV-ABCs or NU-ABCs (120±73, 24±26 or, 2±2, respectively; *P*<0.0001, Figure 2D). In addition, cell proliferation as measured at day 21 by MTT assay was also highly increased in IPF-ABCs compared to HV-ABCs and NU-ABCs (*P*<0.0001, Figure 2E). Based on our scRNAseq data ^16, 22, 23^ we tested whether EGFR ligand production was up-regulated in IPF-ABCs. Indeed, conditioned medium of 3D organoid cultures of IPF-ABCs showed higher levels of amphiregulin (AREG) than HV-ABCs or NU-ABCs (*P*<0.0001), whereas HBEGF or TGF-β were not detectable. Bronchosphere counts correlated with amphiregulin levels at day 14 (*P*<0.0001, r^2^=0.69, Figure 2F).

**Figure 2.**
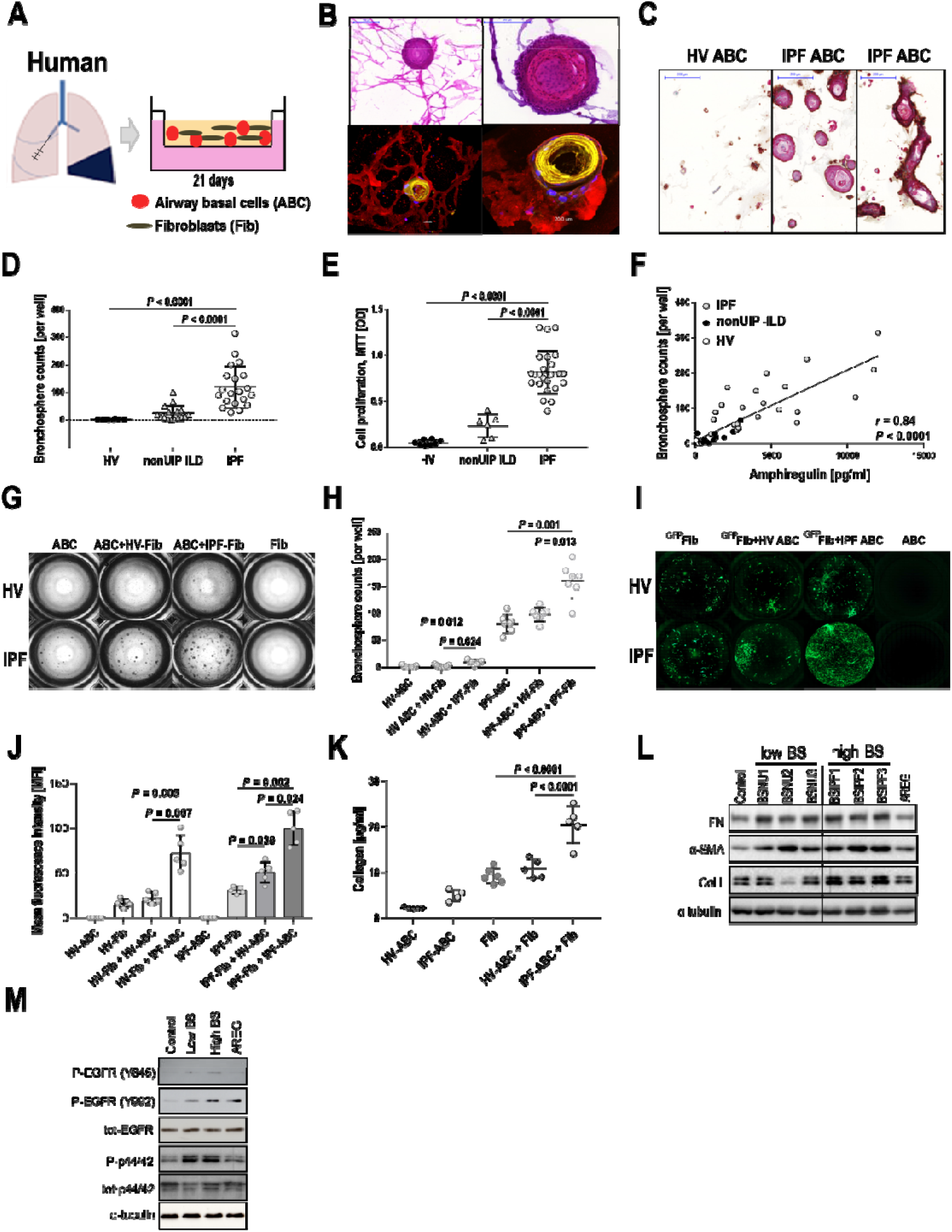
IPF-ABCs generate more bronchospheres and increase fibroblast proliferation and collagen production compared to non-IPF cells in a 3D organoid model. **(A)** Airway basal cells w/wo human lung fibroblasts were cultured in matrigel applying a transwell system. **(B)** IP - ABCs form large spheres which become hollow tube-like structures after 21 days of 3D culture. In the co-culture system of ABCs with lung fibroblasts, fibroblasts surround bronchospheres and form a mesh-like structure. Upper panel: Masson trichrome staining, scale bars 100µm or 200µm as indicated. Lower panel: Confocal immunohistochemistry demonstrates tube formation by IP - ABCs and close interaction with lung fibroblasts (red=vimentin, yellow=CK5/6, blue=TO-PR - 3=nuclei, n=10, scale bar 10µm and 20µm). **(C)** Immuno-histochemisty of evolving bronchospheres stained for CK5/6 in red, p40 in turquoise and beta6 integrin in brown (n=9, scale bars 200µm). **(D)** IPF-ABCs (n=23) generated significantly more and larger spheres than HV-ABCs (n=7; P<0.0001) and NU-ABCs (n=15, P<0.0001). **(E)** Cell proliferation was also significantly increased in IPF-ABCs (n=23) compared to HV-ABCs (n=7, P<0.0001) and NU-ABCs (n=6, P<0.0001) as measured by MTT assay at d21. **(F)** Amphiregulin levels were increased in conditioned medium of bronchospheres from IPF-ABCs (n=23) compared to HV-ABCs (n=7) and NU-ABCs (n=15) and correlated closely with bronchosphere counts. **(G)** Bright field images of raster microscopy of an original experiment (10 independent experiments in triplicate). Lung fibroblasts do not form spheres. Sphere formation by IPF-ABCs is easily detectable. **(H)** In the presence of lung fibroblasts (n=5 IPF-Fib, n=5 HV-Fib), IPF-ABCs and HV-ABCs generate increased numbers of bronchospheres. **(I)** Fibroblast cell lines were transduced with lentiviral vectors encoding GFP. Fibroblast proliferation was highly increased in the presence of IPF-ABCs. **(J)** Mean fluorescence intensity was significantly increased in fibroblast cell lines co-cultured with IPF-ABCs (n=5). **(K)** Collagen levels were detected in conditioned medium a d matrigel by sircol assay at day 63. Collagen production by lung fibroblasts cultured with IPF-ABCs was significantly increased (n=5). **(L)** In addition, fibroblasts cultured in conditioned medium of IPF-ABC derived bronchospheres (high BS) showed an increase in collagen 1A and α- smooth muscle actin (SMA) expression compared to conditioned medium of bronchosphere cultures derived from NU-ABCs (Low BS). **(M)** Lung fibroblasts were treated for 2h with conditioned media of bronchospheres which were harvested at day 14 Pooled conditioned media of

### Presence of fibroblasts increased bronchosphere formation and IPF-ABCs stimulated fibroblast proliferation and collagen production

To assess the interaction of fibroblasts and ABCs we added primary lung fibroblasts to our 3D cell culture system. IPF-ABCs formed bronchospheres similar to the ones observed before, which were now surrounded by a mesh of fibroblasts (Figure 2B). Presence of lung fibroblasts in the 3D cell culture system, particularly primary cells derived from IPF explants, significantly enhanced bronchosphere formation of ABCs (*P*=0.012 and *P*=0.001, Figure 2G and H). Using GFP-transduced fibroblasts, we found that the overall fluorescence was significantly increased by IPF-ABCs in healthy (HV-Fib) as well as IPF fibroblasts (IPF-Fib), (*P*=0.005 and *P*=0.002 respectively, Figure 2I and J), potentially reflecting enhanced proliferation. IPF-ABCs significantly increased collagen production by lung fibroblasts compared to HV-ABCs (*P*<0.0001, Figure 2K). Stimulation of lung fibroblasts with conditioned medium obtained from NU-ABC or IPF-ABC harvested at day 14 induced collagen and alpha-smooth muscle (α-SMA) expression in lung fibroblasts (Figure 2L) as well as induced EGFR phosphorylation (Figure 2M), a finding consistent with significantly increased secretion of amphiregulin by IPF-ABC bronchospheres (Figure 2F).

### Human IPF-ABCs augment bleomycin induced pulmonary fibrosis and induce ultrastructural changes in *RAG2*^*-/-*^ and *NRG* mice

Intratracheal application of human IPF-ABCs or HV-ABCs to uninjured *Rag2*^*-/-*^ mice and NRG mice had no discernible effect (data not shown). Thus, we decided to administer the human IPF-ABCs or HV-ABCs three days after causing the lung minimal injury with a low dose of bleomycin to *RAG2*^*-/-*^ mice (1.2mg/Kg body weight IT, Figure 3A). The administered bleomycin dose resulted in mild fibrotic changes in the lungs (Figure 3B). IPF-ABCs administration highly increased bleomycin induced pulmonary fibrosis compared to HV-ABCs or bleomycin alone as shown by Masson trichrome staining (Figure 3B). Ashcroft score (*P*<0.0001) and hydroxyproline levels (*P*<0.0001) were significantly increased in mice challenged with IPF-ABCs compared to mice challenged with HV-ABCs or bleomycin alone (Figure 3C and D). Similar results were obtained in NOD.Cg-Rag1tm1Mom Il2rgtm1Wjl/SzJZtm (*NRG*) mice (see Figure S2). Masson trichrome staining showed abundant new collagen production centered around the airway structures but also reaching up to the pleura (Figure 3B and F). Starting with day 8 we observed formation of *de novo* airway structures, areas of bronchiolization and formation of cystic structures (Figure 3E and F). Some of the induced lesions resemble pseudoglandular lesions described during lung development or recently described glandular-like epithelial invaginations ^24^ (Figure 3E). As expected, engraftment and proliferation of human ABCs was enhanced in *NRG* mice. In *NRG* mice challenged with IPF-ABCs but not HV-ABCs we regularly observed focal squamous metaplastic lesions of human ABCs (Figure 3G). In some experiments, *NRG* mice were challenged with human IPF-ABCs transfected with a luciferase and GFP encoding vector. Bioluminescence measurements of luciferase expression in mice injected with human IPF-ABCs (day 3-21) showed an increased signal intensity up to day 21 suggesting pulmonary engraftment of human cells and further proliferation (Figure 3H). Introduction of GFP transduced IPF-ABC to the lungs of NOD.Cg-Prkdcscid Il2rgtm1Wjl Tg(CAGGS-VENUS)1/Ztm (*NSG*) venus expressing mice confirmed the engraftment of these cells in the murine lung building up focal squamous metaplastic and pseudoglandular lesions (Figure 3I and Figure S2). Using *NSG* mice we were able to observe engrafted IPF-ABCs adjacent to murine origin areas of glandular-like epithelial invagination structures, supporting an effect of IPF-ABC cells on host resident cells.

**Figure 3.**
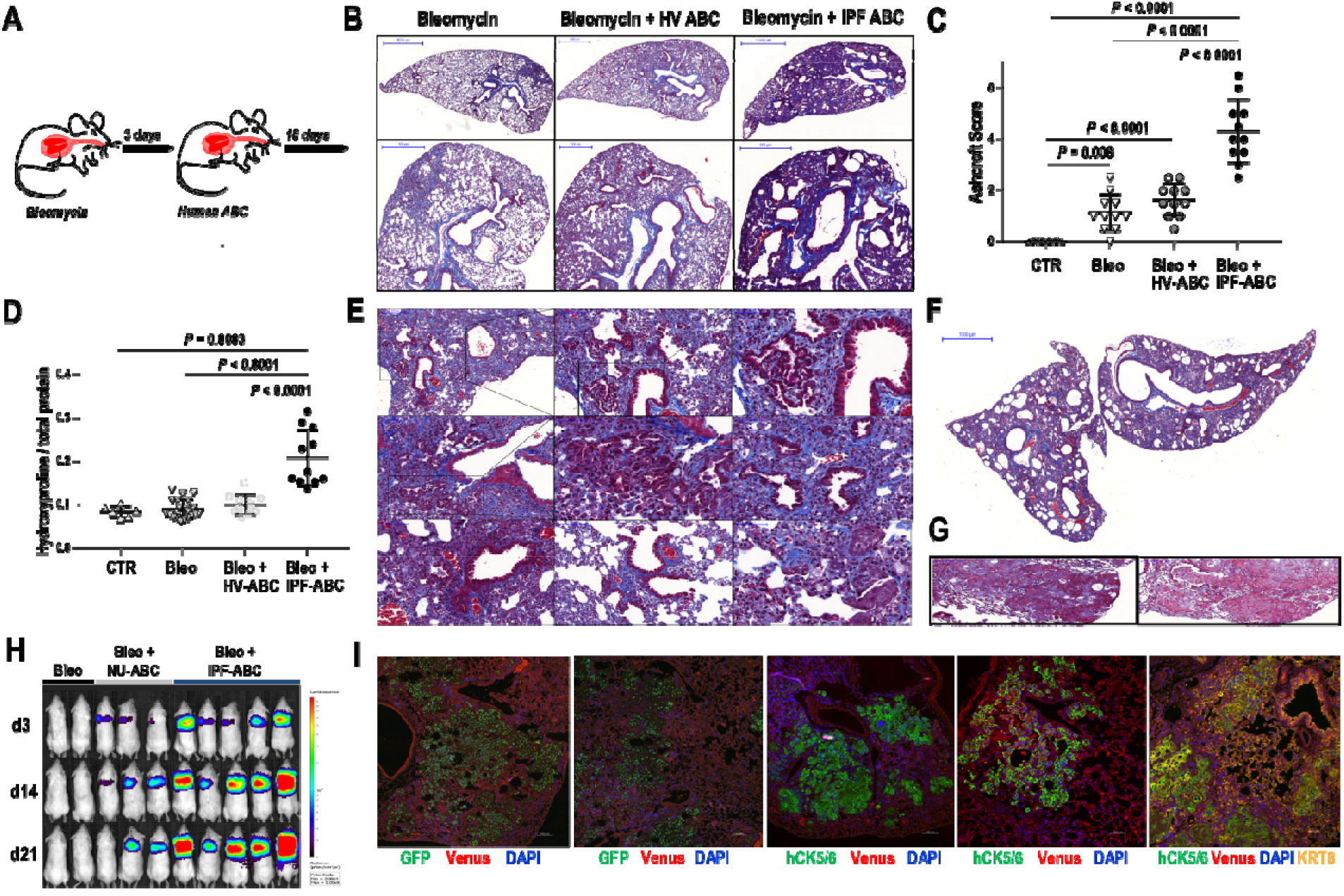
Establishment of a humanized mouse model for IPF based on human ABCs. **(A)** Bleomycin (Bleo) was intratracheally administered to Rag2-/- mice (BL6 background). Three days later human airway basal cells (ABC) derived from either patients with IPF or HV were intratracheally administered. Lungs were harvested at day 21 following Bleo application. **(B)** Representative trichrome stainings of mice challenged with Bleo alone, Bleo and HV-ABCs or Bleo and IPF-ABCs. Fibrosis and cystic lesions are increased in mice challenged with Bleo + IPF-ABCs. **(C-D)** Ashcroft score and hydroxyproline levels were significantly increased in mice challenged with Bleo + IPF-ABCs (n=11) compared to mice challenged with Bleo and HV-ABCs (n=11) or Bleo alone (n=21). **(E**,**F)** Trichrome staining of mice injected with Bleo + IPF-ABCs, shown are representative lesions of 11 different mice. Bronchiolization of the alveolar compartment and de novo generation of airway structures is highly increased in these mice compared to mice challenged with Bleo alone or Bleo + HV-ABCs. These bronchiolar lesions look often bizarre and undirected. Others appear to resemble honeycomb cysts. New collagen synthesis (shown in blue) is seen predominately around the bronchiolar lesions. Rarely structures resembling fibroblast foci could be detected. **(G)** In some experiments with IPF-ABCs in NRG mice metaplastic squamous lesions evolved. **(H)** Mice were challenged with human IPF-ABCs transfected with a luciferase and GFP encoding vector (n=10, 3 replicates). Shown are representative bioluminescence measurements of luciferase expression in control NRG mice (Bleo) and NRG mice injected with transduced human IPF-ABCs and NU-ABCs (day 3-21). **(I)** Engraftment of human IPF-ABCs into venus expressing NSG lungs was detected by confocal microscopy. Human cytokeratin (hCK) 5/6, eGFP, CK8, nuclear DAPI and Venus-expression were detected. Human IPF-ABCs generate focal metaplastic lesions and pseudoglandular lesions in murine lungs. Some of the pseudoglandular lesions are also derived from murine cytokeratin (CK)8+airway epithelial cells. For statistical comparison **(C**,**D)** ANOVA with Tukey correction for multiple testing was used.

### Single cell sequencing of IPF-ABCs and NU-ABCs identifies SRC as a potential therapeutic target

Connectivity MAP analysis using the gene expression profile of IPF-ABC identified a list of substance classes predicted to reverse this IPF-ABC signature: three perturbagen classes had a maximum summary connectivity score of -100: SRC-inhibitors, MEK-inhibitors and loss of function of C2 domain containing protein kinases (Figure 4A) ^25^. Additionally, SRC-inhibition was the second best candidate with a summary connectivity score of -99.94 if we analyzed the ABC signature associated with mortality in a large cohort of IPF patients that we recently published^16^.

**Figure 4.**
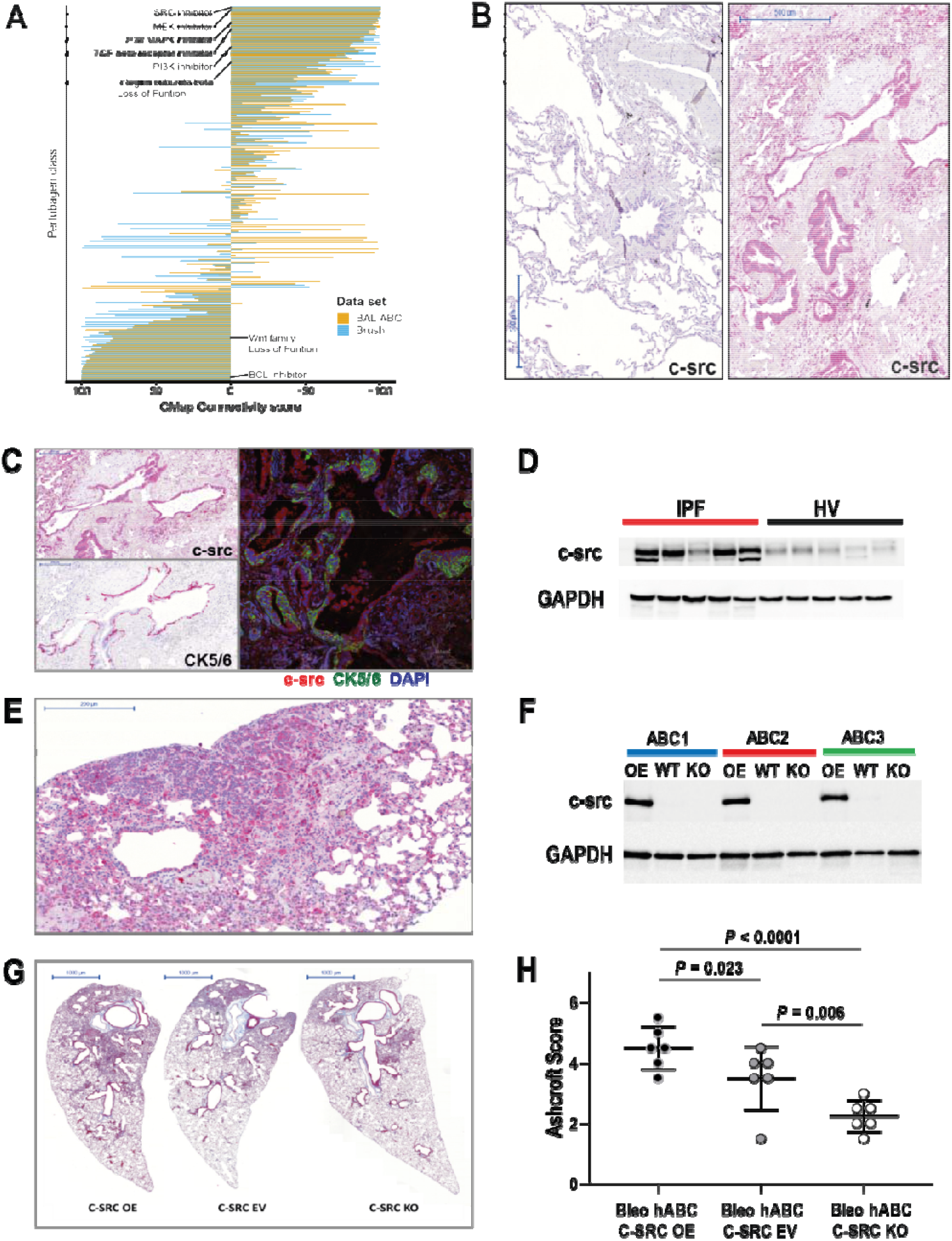
CMap of scRNAseq data identifies potential treatment strategies such as src inhibition. Overexpression of c-src in IPF lungs and cellular effects of c-src over-expression and knockout. **(A)** Bar plots of summary CMap Connectivity scores of the clue.io analysis, dodged by input (ABC mortality signature - yellow, scSeq-derived DEGs IPF vs nonUIP ILD control in ABCs - blue). Inhibition of SRC had the lowest CMap Connectivity score, i.e. potentially reversing the input gene signatures. **(B)** In normal human lung tissue c-src expression was low, only few macrophages were stained positively. In contrast c-src expression of IPF lung tissue was high. Macrophages and epithelial cells highly expressed c-src as well as lymphocytes within lymphoid follicles. **(C)** Most of the epithelial cells expressing c-src were also positive for CK5/6 identifying them as basal cells. **(D)** c-src expression was high in lung tissue homogenates of IPF patients compared to healthy donors. **(E)** Immunohistochemistry of murine lungs of the described humanized mouse model also showed high c-src expression of macrophages and airway epithelial cells. **(F)** C-src expression was either overexpressed, untouched (EV) or knockout in human ABCs using a lentiviral vector and resulting c-src expression was measured using Western blot. **(G**,**H)** C-src overexpression in human IPF-ABCs which were injected into NRG mice lead to increased pulmonary fibrosis, while knockout of c-src downregulated the induced fibrosis (mean ± SD, n=6, each group, 2 replicates). For statistical comparison repeated measures ANOVA with Tukey correction for multiple testing was used.

### SRC expression is increased in lung tissues of IPF patients and murine lungs of the described humanized mouse model

Based on the results of the described *in-silico* analyses, we analyzed SRC protein expression in IPF-tissues and ABCs. SRC was highly increased in IPF lung tissues (Figure 4B), specifically in epithelial cells covering fibroblast foci and within areas of bronchiolization. SRC staining of macrophages was observed in both IPF and control lungs (Figure 4C and Figure S4). SRC was increased in lung homogenates of IPF lung tissues compared to healthy lung tissue (Figure 4D). Impressively, SRC protein expression was also observed in the xenograft IPF-ABC mouse model described above, primarily in areas of aberrant airway generation, bronchiolization and in glandular-like epithelial invagination lesions (Figure 4E). As expected, SRC staining of macrophages was also observed in the mouse lung (Figure 4E).

### SRC overexpression in IPF airway basal cells increased cell invasion and fibrosis, while SRC knock-down attenuated fibrosis

IPF-ABCs were transduced with different vectors which lead to overexpression of SRC, knockdown of SRC or expression of GFP as control (empty vector (EV)). Effects on SRC expression were confirmed by Western blot (Figure 4F). We then tested the effects of modifying SRC expression in IPF-ABCs on fibrosis in the IPF-ABC xenograft model described above. Forced overexpression of SRC in IPF-ABCs (SRC^+^ IPF-ABC) resulted in enhanced fibrosis and cellular remodeling in the alveolar compartment of *NRG* mouse lungs, whereas SRC knockdown in IPF-ABC cells (SRC^-^ IPF-ABC) from the same donor caused markedly reduced remodeling (Figure 4G). Quantification of these results using the Ashcroft score confirmed both, a significant increase in fibrosis in SRC^+^ IPF-ABC and a decrease in fibrosis in SRC^-^ IPF-ABC compared to controls (*P*=0.023, *P*=0.006, respectively Figure 4H).

### The SRC inhibitor saracatinib completely abrogates IPF-ABC bronchosphere formation and attenuates fibroblast proliferation *in-vitro*

Based on the Connectivity MAP predictions, we tested whether saracatinib, a known SRC inhibitor previously tested in human cancer is able to modulate IPF-ABC phenotype ^26^. When IPF-ABC were cultured in 3D and were treated with saracatinib, nintedanib, pirfenidone or vehicle, we observed a complete abrogation of bronchosphere formation at saracatinib concentrations of 600nM, 210nM, and 75nM, whereas nintedanib and pirfenidone had no visible effect on bronchosphere formation (Figure 5A). The results were similar in cells obtained from 12 different individuals with IPF (*P*<0.0001, Figure 5B). Saracatinib did not affect cellular vitality of ABCs in all used concentrations (Figure S5) and these concentrations were considered equivalent to clinically relevant doses ^27^. Similar data were obtained by testing cell proliferation in the MTT assay (*P*<0.0001, Figure 5C). In the co-culture model of IPF-ABC with lung fibroblasts, saracatinib completely blocked bronchosphere formation at the concentrations of 600nM, 210nM and 75nM, had a lower effect at 25nM, and did not have an effect at 8nM (Figure 5D-F). In contrast to the single culture model described above, in the IPF-ABC fibroblast co-culture model, pirfenidone and nintedanib significantly reduced bronchosphere formation, but not completely (Figure 5D-F), potentially reflecting an effect of these drugs on the fibroblast component of this interaction. Fibroblast proliferation in the co-culture was also substantially lower in the presence of saracatinib (Figure 5G and H).

**Figure 5.**
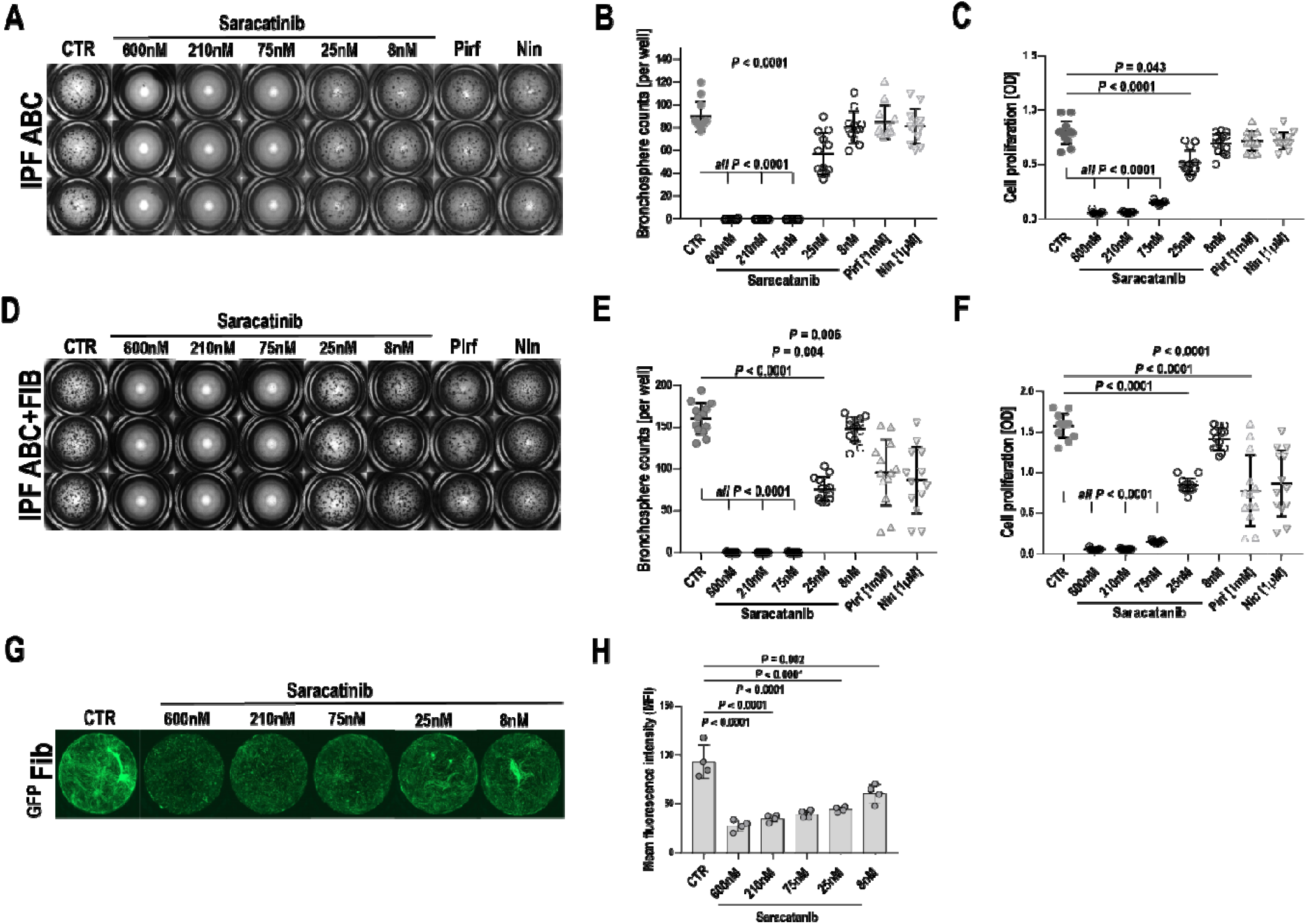
SRC-inhibitor saracatinib abrogates bronchosphere formation. **(A-E)** Bronchosphere assays of human IPF-ABCs were performed w/wo IPF lung fibroblasts for 21 days. Cell cultures (n=12, in triplicates) were treated with either vehicle, saracatinib (600, 210, 75, 25, 8 nM), pirfenidone (1m) or nintedanib (1µM). **(B)** Saracatinib abrogated sphere formation dose-dependently while a dose of pirfenidone or nintedanib, which was higher than us ally applied in humans, did not change bronchosphere counts (mean ± SD). **(C)** Cell proliferation as measured by MTT assay was significantly reduced by saracatinib treatment in a dose dependent manner and slightly by pirfenidone and nintedanib (mean ± SD). **(D**,**E)** Saracatinib treatment also abrogated sphere formation dose-dependently in the presence of IPF lung fibroblasts while pirfenidone and nintedanib reduced significantly, but not abrogated, sphere formation in the presence of fibroblasts (mean ± SD). **(F)** Cell proliferation as measured by the MTT assay was significantly reduced by saracatinib treatment in a dose dependent manner and less reduced by nintedanib or pirfenidone treatment (mean ± SD). **(G**,**H)** In the same model we used GFP+fibroblasts to study fibroblast proliferation. Saracatinib treatment also showed a dose-dependent effect upon fibroblast proliferation (mean ± SD, n=4, 2 replicates). For statistical comparison repeated measures ANOVA with Tukey correction for multiple testing was used.

### Saracatinib attenuated fibrosis and bronchialization *in-vivo*

To test the effect of saracatinib on IPF-ABC induced fibrosis and remodeling *in-vivo*, we returned to the minimal injury xenotransplant model in NRG mice. Overall, we used IPF-ABCs from 38 different individuals with IPF in these experiments. Per IPF subject, IPF-ABCs were used in two animals, one treated with saracatinib and one with vehicle control, to account for interindividual variability. To test the effect of saracatinib on engraftment and development of fibrosis we started treatment at day 4 post injury, one day post IPF-ABC installation for a total of 18 days (Figure 6A). Oropharyngeal saracatinib in a dose of 10mg/kg once daily significantly reduced fibrosis at day 21 as measured by Ashcroft score (4.4±0.9, 1.4±0.7, *P*<0.0001, Figure 6B) and hydroxyproline/total-protein levels (2.4±1.0, 0.7±0.6, P=0.0004, Figure 6C). To test the effect of saracatinib on established fibrosis we performed another set of experiments in which treatment was delayed to day 8 when cell engraftment and fibrosis were already established. Saracatinib treatment again significantly reduced remodeling and fibrosis (Figure 6D) as was reflected by significant reductions in the Ashcroft score (P=0.04, Figure 6E) and hydroxyproline content (P=0.047, Figure 6F).

**Figure 6.**
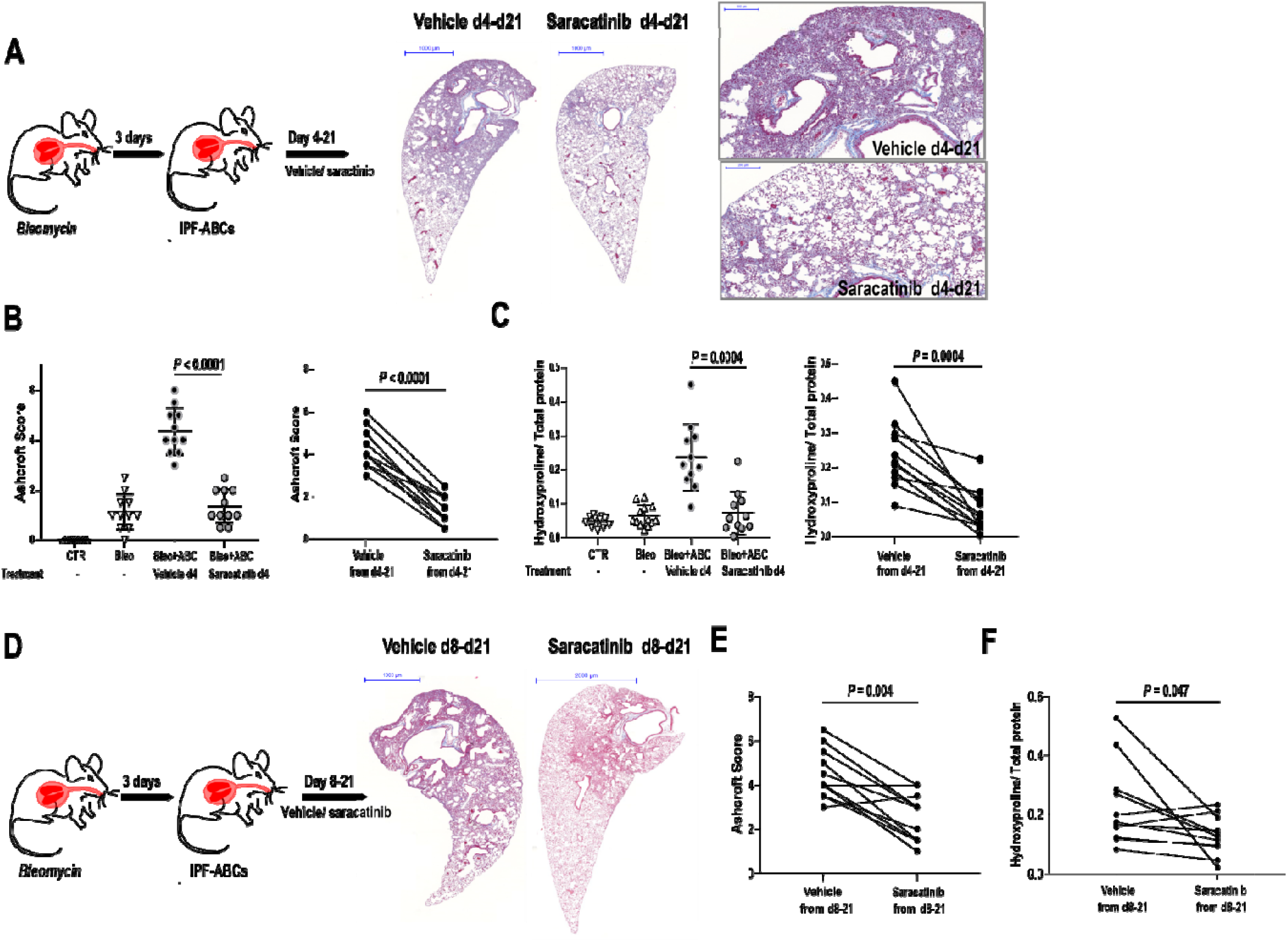
SRC-inhibitor saracatinib attenuates fibrosis in the humanized mouse model. *NRG* mice were intratracheally injected with a low do e of bleomycin and 3 days later with human ABCs derived from IPF patients. Mice (n=11 each group, 4 replicates (all data shown)) were treated oropharyngeally with or without saracatinib in a dose of 10mg/kg once daily starting at day 4 (A-C) or at day 8 (D-F). **(A-C)** Treatment with saracatinib from day 4 significantly reduced the evolution of fibrosis as measured by Ashcroft score (**B**, mean ± SD) and hydroxyproline levels (**C**, mean ± SD). **(D-F)** Saracatinib treatment from day 8 to day 21 also significantly reduced fibrosis but effect was less than with immediate treatment as measured by Ashcroft score **(E)** and hydroxyproline levels **(F)**. All P values were determined by a paired comparison (t-test) testing the effect of saracatinib treatment for each human IPF cell line.

## DISCUSSION

In this paper we demonstrate that airway basal cells which are found in areas of remodeling and bronchiolization and adjacent to fibroblastic foci in the IPF lung have unique properties and are reprogrammed. Transcriptionally, IPF-ABCs are substantially different and exhibit enhanced stemness, ECM sensing and EGF signaling consistent with cellular dedifferentiation. Using a 3D organoid model, we demonstrate that IPF-ABCs give rise to more bronchospheres compared to normal or nonUIP ILD cells. Co-culture experiments with fibroblasts show a close interaction of both cell types which results in augmented bronchosphere formation by IPF-ABC as well as enhanced proliferation and extracellular matrix (ECM) deposition by fibroblasts. Intratracheal application of IPF-ABCs into minimally injured lungs of immunocompromised mice leads to severe fibrosis and remodeling of the alveolar compartment including the evolution of honeycomb cyst-like structures. Connectivity analysis suggested that gene expression changes in IPF-ABCs can be reversed by SRC inhibition and enhanced SRC expression and activity were observed in IPF lungs. Saracatinib, a potent SRC inhibitor, modulated the *in-vitro* and *in-vivo* IPF-ABC induced profibrotic changes.

In this paper we provide the first demonstration that ABCs from patients with IPF are functionally different and have profibrotic effects *in-vivo* and *in-vitro*. ABCs are considered an airway stem cell population, capable of proliferation and self-renewal and can give rise to all types of airway epithelial cells ^11^. Moreover, ABCs have been implicated in COPD, asthma and lung cancer ^11, 28^. In recent years, there have been cumulative evidence that in the IPF lung, ABCs migrate from their airway niche to the lung parenchyma, populating areas of honeycomb cysts and covering fibroblastic foci ^6-8, 10, 16, 29^. However, so far, it was unclear whether ABCs were acting as active players in lung remodelling or whether they served as merely innocent bystanders. We provide several lines of evidence that IPF-ABCs have substantial profibrotic properties and are dedifferentiated. The first clue comes from the observation that unlike HV-ABCs or NU-ABCs, IPF-ABCs generate numerous, well developed and multilayered bronchospheres in the 3D culture model system. This *in-vitro* model is widely used in oncology and developmental biology as an indicator of cellular stemness in formation of tumor spheres and organoids. Our data demonstrate that ABCs of IPF patients differ from those of healthy volunteers by their enhanced capacity to from bronchial like structures in vitro that potentially reflect an exaggerated stem cell phenotype and dedifferentiation of these cells. This is also supported by the scRNAseq results, as IPF-ABCs overexpress known stemness genes; such as KLF4, a transcription factor used for induction of pluripotency and known to be associated with stemness and epithelial-mesenchymal transition (EMT) ^30, 31^; MYC, a transcription factor also used for induction of pluripotency and stemness ^31, 32^; AP1 forming c-JUN and FOSL1 (also known as the proto-oncogene c-fos) which are downstream transcription factors of genes inducing pluripotency ^31, 33, 34^ and important oncogenes for cancer stem cells ^35^. Moreover, PTEN expression was highly decreased in IPF-ABCs and decreased PTEN expression is associated with increased PI3K signaling and increased stem cell renewal. Of interest, the signature of KRT17^high^IPF-ABCs appears to be similar to the recently published signature of KRT17^+^aberrant basaloid cells which lack KRT5 ^17, 18, 21, 36^.

The second line comes from the co-culture experiments that suggested a self-amplifying interaction between ABCs and fibroblasts. The presence of lung fibroblasts, increased evolution of bronchospheres, whereas ABCs stimulated proliferation and collagen production of fibroblasts. In both cases, the effect was stronger when the cells used were obtained from individuals with IPF. Of note, TGF-β was not produced by IPF-ABCs. Our scRNAseq data suggested that one profibrotic factor released by IPF-ABCs is CTGF. Another contributing mechanism to this phenomenon involves the EGFR axis - scRNAseq revealed that amphiregulin expression is significantly increased in IPF-ABCs compared to NU-ABCs and that the bronchospheres generated from IPF-ABCs secrete significantly higher concentrations of amphiregulin compared to both HV-ABCs and NU-ABCs. Conditioned media from IPF-ABC bronchosphere cultures induced phosphorylation of EGFR in fibroblasts. Generally, the role of EGFR ligand family members in fibrosis has been studied mainly in other organs with sometimes conflicting results. However, amphiregulin has been most consistently shown profibrotic effects in models of liver, kidney and heart fibrosis ^37-40^. In the lung, amphiregulin was proposed as a mediator of TGFB1 mediated pulmonary fibrosis in the triple transgenic mouse model ^41^, but without significant follow up. Our results suggest that amphiregulin may be a frequently overlooked major mediator of the interaction of ABCs and fibroblasts in fibrosis.

The third line of evidence comes from the humanized model of lung fibrosis we established. Human IPF-ABCs, but not HV-ABCs, induced abundant bronchiolization, airway enlargement, cyst formation and frequently pleural extending fibrosis, hallmark features of UIP histology missing in commonly used animal models of pulmonary fibrosis. Our model clearly demonstrates that IPF-ABCs induce a vast remodeling and deconstruction of the murine alveolar lung architecture. Taken together these lines of evidence support the unique profibrotic properties of ABCs obtained from lungs of patient with IPF.

The reason why IPF-ABCs are different is unclear, but they are clearly very different from NU-ABCs and show a de-differentiated phenotype. Recent publication call this phenotype hyper- and metaplastic and indeed did we find metaplastic lesions of human basal cells in our mouse model. We speculate that cigarette smoking, exposure to other environmental factors, recurrent airway infections and genetic background may lead to repetitive airway epithelial barrier injury and a chronic wound healing response in IPF, which then may have an impact on ABC gene expression as recently described for asthma and TH2 inflammation ^42^. Genetic risk factor for IPF such as polymorphisms in MUC5B, desmoplakin and AKAP13 are linked to epithelial barrier function and AKAP13 and desmoplakin, unlike MUC5B, are highly expressed by ABCs ^43^. Our scRNAseq data demonstrate increased expression of the stress induced keratins KRT6 and KRT17 ^44^, upregulation of key molecules of epithelial barrier function (claudin 1 and 4), of several integrins which regulate ECM composition (β6, Vα, β1, α6 and α2), markers of epithelial cells senescence such as GDF15 ^45^, and the EGFR ligands AREG and HBEGF in IPF-ABCs. Thus, the transcriptional phenotype of IPF-ABCs may represent the end results of the lung response to repetitive epithelial injuries ^46^, which trigger exaggerated repair processes including dedifferentiation and epithelial-mesenchymal crosstalk ^6, 47^.

Another novel feature of our study is the focus on SRC inhibition in pulmonary fibrosis and especially as mediator of IPF-ABC profibrotic effects. Based on our scRNAseq data and our recently published BAL data set ^16^ we tested for potentially beneficial drug candidates using connectivity map. Our *in-silico* analyses indicated that SRC-inhibitors may be capable of reversing the profibrotic basal cell phenotype which we observe in IPF. SRC is a hub integrating multiple pathways including integrin and receptor kinase signaling ^48, 49^, and results in phosphorylation of various substrates such as STAT3, FAK, JNK, EGR, AKT, and PI3K. Mechanosensitive activation of Beta1 integrin and the downstream signaling via focal adhesion kinase (FAK) and SRC are increased in multiple cancers and linked to proliferation, migration and invasion of cells ^48, 50^. SRC was also shown to align to the EGFR molecule to facilitate it’s signaling both upstream and downstream and is linked to stemness of cancer cells ^48, 51-54^. SRC signaling is closely linked to epithelial injury ^55-62^ induced by various noxious agents including cigarette smoking ^63^. A role for SRC has been proposed in fibrosis in multiple organs but most of this work focused on the role of SRC in fibroblasts ^64-69^. We found abundant SRC expression in ABCs covering the fibroblast foci in IPF lungs. Overexpression of SRC in ABCs resulted in an increase in fibrosis and knockout of SRC reduced fibrosis in our humanized mouse model of IPF. We tested the SRC inhibitor saracatinib (AZD0530) ^70-73^ in our *in-vitro* and *in-vivo* models. Saracatinib completely abrogated sphere formation in the 3D organoid model. In the co-culture model, saracatinib had an effect on both sphere formation and fibroblast proliferation suggesting that SRC inhibition by saracatinib may disrupt the profibrotic HV-ABCs crosstalk between IPF-ABCs and fibroblasts. Similar effects were also observed in our humanized mouse model, where saracatinib substantially attenuated IPF-ABC induced pulmonary fibrosis, bronchiolization and cyst formation. This effect was also observed when saracatinib was administered during established fibrosis. Taken together, these findings suggest that saracatinib treatment efficiently blocked human IPF-ABC engraftment and proliferation.

In conclusion, our data indicate a dedifferentiated phenotype and profibrotic role of ABCs in IPF. In recent years there has been a significant increase in the understanding of the mechanisms of pulmonary fibrosis ^1, 2^. Roles for distinct lung cell populations such as fibroblasts and myofibroblasts, macrophages and even alveolar epithelial cells have been proposed. While, ABCs have been noticed in the IPF lung, mechanistic studies evaluating their properties, and role in pulmonary fibrosis have been missing. Our results provide clear evidence that human IPF-ABCs are dedifferentiated and have profibrotic properties *in-vitro* and *in-vivo*, and that interventions that address their properties reverse fibrosis. These findings position the ABC as a key cell in the pathogenesis of human pulmonary fibrosis and thus a novel cellular target for therapeutic interventions.

## MATERIALS AND METHODS

A comprehensive description of all methods is available in the supplementary information.

### Study population

For the described experiments in total bronchial brushes from 68 patients with IPF, 25 patients with nonUIP fibrotic ILD and 18 healthy volunteers of an older age (>50 years) were obtained. IPF diagnosis was established by a multidisciplinary board according to the ATS/ERS criteria ^3, 74^. The study was approved by local ethics committees and all included individuals signed informed consent. For more details see supplement.

### Immunohistochemistry

Immunohistochemistry of lung tissues was performed using a standard protocol and as recently described ^16^. Detailed information is given in the supplement.

### Isolation of airway basal cells

Airway basal cells were isolated from bronchial brushes of sub-segmental bronchi of the right lower lobe as recently described ^75^. Detailed information regarding cell isolation is given in the supplement.

### Human 3D bronchosphere assay

ABCs, either derived from IPF patients or HV, were cultured in matrigel (Corning) in a transwell system with or without lung fibroblasts either derived from patients with IPF or normal lung for 21 days (if not otherwise indicated) in the incubator (5% CO_2_, 37°C). Medium exchange (BEGM (Lonza, Basel, Switzerland, #CC-3170)/ DMEM (Dulbecco’s Modified Eagle Medium (Gibco, Swit Fisher Scientific/Germany); ratio 1:1) was done every 7 days. In some experiments lentiviral transduced GFP expressing fibroblasts were used. Conditioned medium of the 3D organoid culture (day 7 and day 14) was used for ELISA and fibroblast stimulation experiments. Sircol assay (Scientific-Biocolor/UK, #S1000) was performed as recommended by the manufacturer on matrigel and conditioned medium harvested after 6 weeks of cell culture. Bronchosphere formation was documented by Axio Vert.A1/Zeiss/Germany and Axio Observer Z1/Zeiss/Germany. For histology, immunohistology and confocal laser microscopy 3D cell cultures were cryopreserved using Tissue-Tek and cryomolds (Hartenstein/Germany; #CMM). Detailed information is given in the supplement.

### Single cell RNA sequencing of brush cells

Single cell sequencing libraries of brush cells from nine IPF patients and six non UIP ILD control patients were generated as previously described ^17^ using the 10x Genomics Chromium Single Cell 3’ v2 kits according the manufacturer’s instructions. Libraries were pooled and sequenced on an Illumina HiSeq 4000 aiming for 150 million reads per library. zUMIs pipeline was used for subsequent processing of reads. Data analysis and visualization were performed using the R package Seurat.

## Data availability

Sequencing data will be available on the GEO repository. Gene expression of ABCs of IPF and nonUIP ILD controls were compared using a Wilcoxon Rank Sum test with Bonferroni correction of p-values for multiple testing.

### Connectivity Map (CMap) analysis

Broad Institute’s CLUE platform (https://clue.io) was used to identify a potential molecular mechanism of action which could reverse A) the deviating gene expression profile of ABCs in IPF as identified by scSeq in this study and B) the ABC gene expression signature associated with mortality in IPF by bulk RNA sequencing as recently described by us ^16^. Pertubagen classes were extracted and visualized as bar plots of the summary connectivity scores split by the origin of the input gene profiles.

### Humanized pulmonary fibrosis model with intratracheal administration of human ABCs into mice *in-vivo*

Different mice strains were used for the experiments as indicated. B6;129Sv-Rag2tm1Fwa/ZTM^76^ (*Rag2*^*-/-*^) and NOD.Cg-Rag1tm1Mom Il2rgtm1Wjl/SzJZtm^77^ (*NRG*) were obtained from the central animal facility (Hannover Medical School, Hannover). NOD.Cg-Prkdcscid Il2rgtm1Wjl Tg(CAGGS-VENUS)1/Ztm (*Venus-NSG*) mice were kindly provided by Dr. Wiebke Garrels (Hannover Medical School). Mice received bleomycin dissolved in sterile saline at the dose of 1.2mg/kg intratracheally on day 0. IPF-ABCs, HV-ABCs (*Rag 2*^*-/-*^: 0.3×10^5^, *NRG* and *NSG*: 0.2×10^5^) were injected intratracheally on day 3. In some experiments human IPF-ABCs transduced with a lentiviral vector encoding for GFP and luciferase were injected. Pairs of mice received the same IPF-ABC line and were later treated either with vehicle or saractinib (kindly provided by Leslie Cousens AstraZeneca) in a dose of 10mg/kg, i.g. once daily. For histological and immunohistological analysis, the trachea was cannulated and lungs were insufflated with 4% paraformaldehyde in PBS at a pressure of 25 cm H_2_O, followed by removal of the heart and inflated lungs en bloc and immersion in 4% paraformaldehyde overnight at 4°C, and the tissues were processed to paraffin wax or were cryopreserved using Tissue Tek® O.C.T.TM compound. H&E, Masson trichrome stains and Ashcroft score were performed according to a standard protocol ^78, 79^. Collagen content was determined by quantifying hydroxyproline levels according to manufacturer’s description. For further details, see supplementary information.

## Acknowledgment

The authors thank Stefanie Reuss, Victoria Wirtz and Angelina Malassa for their technical support.

## Authors contributions

### Conceptualization

Prasse, Jäger, Schupp, Kaminski

### Methodology

Prasse, Jäger, Schupp, Nazarenko, Wygrecka, Schambach, Zweigerdt, Kaminski

### Investigation

Schupp, Jaeger, Plappert, Terwolbeck, Adams, Kayser, Jonigk, Prasse, Klatt, Wygrecka, Lienenklaus, Garrels, Engelhard

### Formal Analysis

Prasse, Schupp, Kaminski, Jäger, Kayser, Wygrecka, Schambach, Jonigk, Nazarenko, Lienenklaus

### Statistical Analysis

Schupp, Adams, Prasse, Jäger, Wygrecka

### Writing-original Draft

Prasse, Schupp, Kaminski

### Writing-Review & Editing

Schupp, Jaeger, Plappert, Terwolbeck, Adams, Kayser, Jonigk, Schambach, Klatt, Wygrecka, Nazarenko, Kempf, Lienenklaus, Garrels, Engelhard, Zweigerdt, Kaminski, Prasse

### Resources

Prasse, Kayser, Jonigk, Schupp, Kaminski, Schambach, Garrels, Lienenklaus, Zweigerdt, Nazarenko

### Supervision

Prasse, Schupp, Kaminski

### Funding Acquisition

Prasse, Kaminski, Schupp

## REFERENCES

1. Lederer, D.J. & Martinez, F.J. Idiopathic Pulmonary Fibrosis. N Engl J Med 378, 1811–1823 (2018).

2. King, T.E., Jr., Pardo, A. & Selman, M. Idiopathic pulmonary fibrosis. Lancet 378, 1949–1961 (2011).

3. Raghu, G. et al. Diagnosis of Idiopathic Pulmonary Fibrosis. An Official ATS/ERS/JRS/ALAT Clinical Practice Guideline. Am J Respir Crit Care Med 198, e44–e68 (2018).

4. Verleden, S.E. et al. Small airways pathology in idiopathic pulmonary fibrosis: a retrospective cohort study. Lancet Respir Med 8, 573–584 (2020).

5. Chilosi, M. et al. Abnormal re-epithelialization and lung remodeling in idiopathic pulmonary fibrosis: the role of deltaN-p63. Lab Invest 82, 1335–1345 (2002).

6. Jonsdottir, H.R. et al. Basal cells of the human airways acquire mesenchymal traits in idiopathic pulmonary fibrosis and in culture. Lab Invest 95, 1418–1428 (2015).

7. Smirnova, N.F. et al. Detection and quantification of epithelial progenitor cell populations in human healthy and IPF lungs. Respir Res 17, 83 (2016).

8. Xu, Y. et al. Single-cell RNA sequencing identifies diverse roles of epithelial cells in idiopathic pulmonary fibrosis. JCI Insight 1, e90558 (2016).

9. Vaughan, A.E. et al. Lineage-negative progenitors mobilize to regenerate lung epithelium after major injury. Nature (2014).

10. Plantier, L. et al. Ectopic respiratory epithelial cell differentiation in bronchiolised distal airspaces in idiopathic pulmonary fibrosis. Thorax 66, 651–657 (2011).

11. Rock, J.R., Randell, S.H. & Hogan, B.L. Airway basal stem cells: a perspective on their roles in epithelial homeostasis and remodeling. Dis Model Mech 3, 545–556 (2010).

12. Rock, J.R. et al. Basal cells as stem cells of the mouse trachea and human airway epithelium. Proc Natl Acad Sci U S A 106, 12771–12775 (2009).

13. Ghosh, M. et al. Exhaustion of Airway Basal Progenitor Cells in Early and Established Chronic Obstructive Pulmonary Disease. Am J Respir Crit Care Med 197, 885–896 (2018).

14. Zuo, W.L. et al. EGF-Amphiregulin Interplay in Airway Stem/Progenitor Cells Links the Pathogenesis of Smoking-Induced Lesions in the Human Airway Epithelium. Stem Cells 35, 824–837 (2017).

15. Fukui, T. et al. Lung adenocarcinoma subtypes based on expression of human airway basal cell genes. Eur Respir J 42, 1332–1344 (2013).

16. Prasse, A. et al. BAL Cell Gene Expression Is Indicative of Outcome and Airway Basal Cell Involvement in Idiopathic Pulmonary Fibrosis. Am J Respir Crit Care Med 199, 622–630 (2019).

17. Adams, T.S. et al. Single-cell RNA-seq reveals ectopic and aberrant lung-resident cell populations in idiopathic pulmonary fibrosis. Sci Adv 6, eaba1983 (2020).

18. Habermann, A.C. et al. Single-cell RNA sequencing reveals profibrotic roles of distinct epithelial and mesenchymal lineages in pulmonary fibrosis. Sci Adv 6, eaba1972 (2020).

19. Hosur, V., Farley, M.L., Burzenski, L.M., Shultz, L.D. & Wiles, M.V. ADAM17 is essential for ectodomain shedding of the EGF-receptor ligand amphiregulin. FEBS Open Bio 8, 702–710 (2018).

20. Wang, X. et al. A Disintegrin and A Metalloproteinase-9 (ADAM9): A Novel Proteinase Culprit with Multifarious Contributions to COPD. Am J Respir Crit Care Med (2018).

21. Kobayashi, Y. et al. Persistence of a regeneration-associated, transitional alveolar epithelial cell state in pulmonary fibrosis. Nat Cell Biol 22, 934–946 (2020).

22. Brechbuhl, H.M., Li, B., Smith, R.W. & Reynolds, S.D. Epidermal growth factor receptor activity is necessary for mouse basal cell proliferation. Am J Physiol Lung Cell Mol Physiol 307, L800–810 (2014).

23. Shaykhiev, R. et al. EGF shifts human airway basal cell fate toward a smoking-associated airway epithelial phenotype. Proc Natl Acad Sci U S A 110, 12102–12107 (2013).

24. Aros, C.J. et al. Distinct Spatiotemporally Dynamic Wnt-Secreting Niches Regulate Proximal Airway Regeneration and Aging. Cell Stem Cell (2020).

25. Madala, S.K. et al. MEK-ERK pathway modulation ameliorates pulmonary fibrosis associated with epidermal growth factor receptor activation. Am J Respir Cell Mol Biol 46, 380–388 (2012).

26. Puls, L.N., Eadens, M. & Messersmith, W. Current status of SRC inhibitors in solid tumor malignancies. Oncologist 16, 566–578 (2011).

27. Hu, M. et al. Therapeutic targeting of SRC kinase in myofibroblast differentiation and pulmonary fibrosis. J Pharmacol Exp Ther 351, 87–95 (2014).

28. Ryan, D.M. et al. Smoking dysregulates the human airway basal cell transcriptome at COPD risk locus 19q13.2. PLoS One 9, e88051 (2014).

29. Ambrosini, V. et al. Acute exacerbation of idiopathic pulmonary fibrosis: report of a series. Eur Respir J 22, 821–826 (2003).

30. Ghaleb, A.M. & Yang, V.W. Kruppel-like factor 4 (KLF4): What we currently know. Gene 611, 27–37 (2017).

31. Sridharan, R. et al. Role of the murine reprogramming factors in the induction of pluripotency. Cell 136, 364–377 (2009).

32. Yoshida, G.J. Emerging roles of Myc in stem cell biology and novel tumor therapies. J Exp Clin Cancer Res 37, 173 (2018).

33. Ibrahim, E.E. et al. Embryonic NANOG activity defines colorectal cancer stem cells and modulates through AP1-and TCF-dependent mechanisms. Stem Cells 30, 2076–2087 (2012).

34. Zhou, Y. et al. Endogenous authentic OCT4A proteins directly regulate FOS/AP-1 transcription in somatic cancer cells. Cell Death Dis 9, 585 (2018).

35. Lai, D. et al. PP2A inhibition sensitizes cancer stem cells to ABL tyrosine kinase inhibitors in BCR-ABL(+) human leukemia. Sci Transl Med 10 (2018).

36. Strunz, M. et al. Alveolar regeneration through a Krt8+ transitional stem cell state that persists in human lung fibrosis. Nat Commun 11, 3559 (2020).

37. Perugorria, M.J. et al. The epidermal growth factor receptor ligand amphiregulin participates in the development of mouse liver fibrosis. Hepatology 48, 1251–1261 (2008).

38. Cuevas, M.J., Tieppo, J., Marroni, N.P., Tunon, M.J. & Gonzalez-Gallego, J. Suppression of amphiregulin/epidermal growth factor receptor signals contributes to the protective effects of quercetin in cirrhotic rats. J Nutr 141, 1299–1305 (2011).

39. Liu, L. et al. Amphiregulin enhances cardiac fibrosis and aggravates cardiac dysfunction in mice with experimental myocardial infarction partly through activating EGFR-dependent pathway. Basic Res Cardiol 113, 12 (2018).

40. Rayego-Mateos, S. et al. Role of Epidermal Growth Factor Receptor (EGFR) and Its Ligands in Kidney Inflammation and Damage. Mediators Inflamm 2018, 8739473 (2018).

41. Zhou, Y. et al. Amphiregulin, an epidermal growth factor receptor ligand, plays an essential role in the pathogenesis of transforming growth factor-beta-induced pulmonary fibrosis. J Biol Chem 287, 41991–42000 (2012).

42. Ordovas-Montanes, J. et al. Allergic inflammatory memory in human respiratory epithelial progenitor cells. Nature 560, 649–654 (2018).

43. Allen, R.J. et al. Genetic variants associated with susceptibility to idiopathic pulmonary fibrosis in people of European ancestry: a genome-wide association study. Lancet Respir Med 5, 869–880 (2017).

44. Zhussupbekova, S. et al. A Mouse Model of Hyperproliferative Human Epithelium Validated by Keratin Profiling Shows an Aberrant Cytoskeletal Response to Injury. EBioMedicine 9, 314–323 (2016).

45. Zhang, Y. et al. GDF15 is an epithelial-derived biomarker of idiopathic pulmonary fibrosis. Am J Physiol Lung Cell Mol Physiol 317, L510–L521 (2019).

46. Wang, F., Chen, S., Liu, H.B., Parent, C.A. & Coulombe, P.A. Keratin 6 regulates collective keratinocyte migration by altering cell-cell and cell-matrix adhesion. J Cell Biol 217, 4314–4330 (2018).

47. Volckaert, T. et al. Fgf10-Hippo Epithelial-Mesenchymal Crosstalk Maintains and Recruits Lung Basal Stem Cells. Dev Cell 43, 48–59 e45 (2017).

48. Irby, R.B. & Yeatman, T.J. Role of Src expression and activation in human cancer. Oncogene 19, 5636–5642 (2000).

49. Zhao, J. & Guan, J.L. Signal transduction by focal adhesion kinase in cancer. Cancer Metastasis Rev 28, 35–49 (2009).

50. Matsumoto, T. et al. Targeted expression of c-Src in epidermal basal cells leads to enhanced skin tumor promotion, malignant progression, and metastasis. Cancer Res 63, 4819–4828 (2003).

51. Marquardt, J.U. et al. Human hepatic cancer stem cells are characterized by common stemness traits and diverse oncogenic pathways. Hepatology 54, 1031–1042 (2011).

52. Su, Y.J. et al. Polarized cell migration induces cancer type-specific CD133/integrin/Src/Akt/GSK3beta/beta-catenin signaling required for maintenance of cancer stem cell properties. Oncotarget 6, 38029–38045 (2015).

53. Sun, Q., Wang, Y. & Desgrosellier, J.S. Combined Bcl-2/Src inhibition synergize to deplete stem-like breast cancer cells. Cancer Lett 457, 40–46 (2019).

54. Sham, D., Wesley, U.V., Hristova, M. & van der Vliet, A. ATP-mediated transactivation of the epidermal growth factor receptor in airway epithelial cells involves DUOX1-dependent oxidation of Src and ADAM17. PLoS One 8, e54391 (2013).

55. Severgnini, M. et al. Inhibition of the Src and Jak kinases protects against lipopolysaccharide-induced acute lung injury. Am J Respir Crit Care Med 171, 858–867 (2005).

56. Zhao, T. et al. Role of the PKCalpha-c-Src tyrosine kinase pathway in the mediation of p120-catenin degradation in ventilator-induced lung injury. Respirology 21, 1404–1410 (2016).

57. Li, J. et al. Heat-Induced Epithelial Barrier Dysfunction Occurs via C-Src Kinase and P120ctn Expression Regulation in the Lungs. Cell Physiol Biochem 48, 237–250 (2018).

58. Taniguchi, K. et al. A gp130-Src-YAP module links inflammation to epithelial regeneration. Nature 519, 57–62 (2015).

59. Xiong, C. et al. Pharmacological inhibition of Src kinase protects against acute kidney injury in a murine model of renal ischemia/reperfusion. Oncotarget 8, 31238–31253 (2017).

60. Li, L.F. et al. Mechanical ventilation augments bleomycin-induced epithelial-mesenchymal transition through the Src pathway. Lab Invest 94, 1017–1029 (2014).

61. Oyaizu, T. et al. Src tyrosine kinase inhibition prevents pulmonary ischemia-reperfusion-induced acute lung injury. Intensive Care Med 38, 894–905 (2012).

62. Shan, X. et al. Inhibition of epidermal growth factor receptor attenuates LPS-induced inflammation and acute lung injury in rats. Oncotarget 8, 26648–26661 (2017).

63. Geraghty, P., Hardigan, A. & Foronjy, R.F. Cigarette smoke activates the protooncogene c-src to promote airway inflammation and lung tissue destruction. Am J Respir Cell Mol Biol 50, 559–570 (2014).

64. Gortzen, J. et al. Interplay of Matrix Stiffness and c-SRC in Hepatic Fibrosis. Front Physiol 6, 359 (2015).

65. Ramachandran, S. et al. Hepatitis C virus induced miR200c down modulates FAP-1, a negative regulator of Src signaling and promotes hepatic fibrosis. PLoS One 8, e70744 (2013).

66. Wang, J. et al. Targeting Src attenuates peritoneal fibrosis and inhibits the epithelial to mesenchymal transition. Oncotarget 8, 83872–83889 (2017).

67. Lu, Y.Y. et al. Interaction of Src and Alpha-V Integrin Regulates Fibroblast Migration and Modulates Lung Fibrosis in A Preclinical Model of Lung Fibrosis. Sci Rep 7, 46357 (2017).

68. Skhirtladze, C. et al. Src kinases in systemic sclerosis: central roles in fibroblast activation and in skin fibrosis. Arthritis Rheum 58, 1475–1484 (2008).

69. Yan, Y. et al. Src inhibition blocks renal interstitial fibroblast activation and ameliorates renal fibrosis. Kidney Int 89, 68–81 (2016).

70. Roskoski, R., Jr. Src protein-tyrosine kinase structure, mechanism, and small molecule inhibitors. Pharmacol Res 94, 9–25 (2015).

71. Posadas, E.M. et al. Saracatinib as a metastasis inhibitor in metastatic castration-resistant prostate cancer: A University of Chicago Phase 2 Consortium and DOD/PCF Prostate Cancer Clinical Trials Consortium Study. Prostate 76, 286–293 (2016).

72. Gucalp, A. et al. Phase II trial of saracatinib (AZD0530), an oral SRC-inhibitor for the treatment of patients with hormone receptor-negative metastatic breast cancer. Clin Breast Cancer 11, 306–311 (2011).

73. Molina, J.R. et al. A phase II trial of the Src-kinase inhibitor saracatinib after four cycles of chemotherapy for patients with extensive stage small cell lung cancer: NCCTG trial N-0621. Lung Cancer 85, 245–250 (2014).

74. American Thoracic Society. Idiopathic pulmonary fibrosis: diagnosis and treatment. International consensus statement. American Thoracic Society (ATS), and the European Respiratory Society (ERS). Am J Respir Crit Care Med 161, 646–664 (2000).

75. Hackett, N.R. et al. The human airway epithelial basal cell transcriptome. PLoS One 6, e18378 (2011).

76. Shinkai, Y. et al. RAG-2-deficient mice lack mature lymphocytes owing to inability to initiate V(D)J rearrangement. Cell 68, 855–867 (1992).

77. Pearson, T. et al. Non-obese diabetic-recombination activating gene-1 (NOD-Rag1 null) interleukin (IL)-2 receptor common gamma chain (IL2r gamma null) null mice: a radioresistant model for human lymphohaematopoietic engraftment. Clinical and experimental immunology 154, 270–284 (2008).

78. Huleihel, L. et al. Modified mesenchymal stem cells using miRNA transduction alter lung injury in a bleomycin model. Am J Physiol Lung Cell Mol Physiol 313, L92–L103 (2017).

79. Ashcroft, T., Simpson, J.M. & Timbrell, V. Simple method of estimating severity of pulmonary fibrosis on a numerical scale. J Clin Pathol 41, 467–470 (1988).

