## Supplement for "Airway Basal Cells show a dedifferentiated KRT17^high^Phenotype and promote Fibrosis in Idiopathic Pulmonary Fibrosis"

**Supplementary Methods**

**Experimental Model and Subject Details**

***Human Specimens***

All experiments with human tissue samples were performed under protocols approved by the Institutional Review Boards at Hannover Medical School and University Medical Center Freiburg. All patients and healthy volunteers signed informed consent prior to inclusion to the study. In total, bronchial brushes from 68 patients with idiopathic pulmonary fibrosis (IPF), 25 patients with nonUIP fibrotic ILD and 18 healthy volunteers of an older age (>50 years) obtained. Only patients with an idiopathic UIP and HRCT consistent with a “definite” UIP pattern were included. We did not notice any sex-dependent effects. In addition, lung explant tissues of in total 20 patients with IPF, 10 patients with COPD and normal tissue from 5 donor lungs were used for fibroblast isolation, lung homogenates and immunohistochemistry. IPF diagnosis and other ILD diagnosis was established by a multidisciplinary board according to the American Thoracic Society/ European Respiratory Society criteria ^1, 2^ and was later determined to be consistent with recent guidelines ^3^.

***Mice***

All mouse procedures were conducted in accordance in with the German law for animal protection and the European Directive 2010/63/EU and were approved by the respective local government (Regierungspräsidium Freiburg, Germany; AZ: 35-9185.81/G-14/17) and Lower Saxony State Office for Consumer Protection and Food Safety in Oldenburg/Germany (LAVES); AZ: 33.12-42502-04-15/1896 and AZ: 33.19-42502-04-15/2017), AZ:33.19-42502-04-17/2612). Different mice strains were used for the experiments as indicated. B6;129Sv-Rag2tm1Fwa/ZTM ^4^ (***Rag2^-/-^***) and NOD.Cg-Rag1tm1Mom Il2rgtm1Wjl/SzJZtm ^5^ (***NRG***) were obtained from the central animal facility (Hannover Medical School, Hannover). NOD.Cg-Prkdcscid Il2rgtm1Wjl Tg(CAGGS-VENUS)1/Ztm (***Venus-NSG***) mice were kindly provided by Dr. Wiebke Garrels (Hannover Medical School). *Venus-NSG* were generated with cytoplasmic plasmid injection of a non-autonomous transposon system ^6, 7^. All cells of this mice stock are expressing a native live imaging fluorescent protein Venus ^8^. Mice were housed at the animal unit of the University Medical Center Freiburg or Hannover Medical School (Central Animal Facility). At both institutions mice were housed in specific pathogen free (SPF) conditions in accordance with institutional guidelines and ethical regulations. All animals used in this study had no previous history of experimentation and were naive at the time of analysis. *RAG2^-/-^* mice used in this study were maintained on the C57BL/6J background for >6 generations. All *RAG2^-/-^, NRG* and *Venus-NSG* mice used in this study were male and 8-12 weeks old.

**Method Details**

***Bronchoscopy***

Bronchial epithelial cells were harvested by bronchial brushes of sub-segmental bronchi of the right lower lobe during flexible bronchoscopy. In patients this was done within the routine diagnostic work-up at initial diagnosis. None of the patients received antifibrotic treatment prior to bronchoscopy. Healthy volunteers underwent bronchoscopy voluntarily for study purpose. None of the included subjects was currently smoking.

***Isolation of airway basal cells (ABCs)***

ABCs were isolated from bronchial brushes of sub-segmental bronchi of the right lower lobe using a similar protocol as recently described.^9^ Bronchial brushes were placed in 2 ml of pre-warmed (37°C) Clonetics™ Bronchial Epithelium Cell Growth Medium (BEGM) (Lonza, #CC-3170). Cells were detached from the brush by flicking the brush. Then, airway epithelial cells were pelleted by centrifugation (250×g, 5 min) and disaggregated by resuspension in trypsin/EDTA-solution (0.05 %/0.02 %) (Merck, #L2143) for 5 min at 37°C. Trypsinization was stopped by addition of HEPES buffered saline (Merck, #391340) supplemented with 15% fetal bovine serum (Merck, #TMS-016-B), then cells were again pelleted at 250×g, 5 min. The pellet was washed once with 5 ml of phosphate buffered saline (PBS, Thermo Fisher, #14190250). Afterwards, the cell pellet was resuspended in 5 ml of BEGM and seeded in T25 flasks (Merck, #CLS3056) in BEGM, supplemented with growth factors according to the manufacturer’s instructions. The antibiotics supplied by the manufacturer of BEGM were used and additionally 0.1% amphotericin B (BioWhittaker Amphotericin B Antifungal Lonza, #17-836R) and 1% penicillin-streptomycin (10000 U/ml), (Merck, #A 2212) were added. Cultures were maintained in a humidified atmosphere of 5% CO_2_ at 37°C. Unattached cells were removed by changing of media after 18 hours. Thereafter, media was changed every 7 days and cells were harvested at day 21, when the cells were 90% confluent. Therefore, cells were trypsinized, harvested and counted. Purity of the ABC population was determined by immunocytology (cytokeratin 5/6 staining) of cytospins and always exceeded 98%.

***Isolation of primary fibroblasts***

Fibroblasts were isolated from fresh tissue blocks (0.3cm x 0.3cm x 0.3cm) derived either from IPF lung explants (n=10) or healthy donor lung (n=5), using a protocol previously described by us.^10^ In brief, small blocks of lung tissues were washed three times in PBS at 4°C. Tissue blocks were then cultured in Gibco DMEM (Dulbecco's Modified Eagle Medium, Thermo Fisher Scientific, #11995-065) supplemented with 10% FBS, 1% penicillin/ streptomycin and 0.1% amphotericin B in 6 well plates and resulted in the outgrowth of fibroblasts. Blocks were removed after 12 days. Outgrown fibroblasts were harvested at day 21 when cells reached confluency. Afterwards fibroblasts were trypsinized for 3 min at 37°C, then immediately suspended in DMEM supplemented with 10% FBS, then pelleted at 250×g for 5 min and seeded in T75 flasks (Merck, #CLS3276). Fibroblasts were cultured in DMEM with 10% FBS for 10 days until cells reached confluency, then harvested using the same trypsinization protocol as described above. Fibroblasts were archived in liquid nitrogen in aliquots of 1 x 10^6^ cells per vial in 20% dimethyl sulfoxide (Merck, #276855) + 80% FBS containing DMEM freezing medium for subsequent experiments.

***Bronchosphere assay (3D organoids)***

In order to test sphere formation and evolution of 3D organoids, a protocol described for tumor sphere formation was used ^11^. Human ABCs (10^4^ cells) and/ or primary human lung fibroblasts (10^4^ cells) were added to 50 µl ice-cold matrigel (corning^®^ matrigel^®^ matrix (Corning^®^, #356231) in transwell inserts (Corning lifesciences Costar, #3470) and cultured for 30 min in the incubator (5% CO_2_, 37°C) until the matrigel became stiff. Then 600 µl of a 1:1 ratio of BEGM and DMEM was added below the insert and additional 100 µl on top of the insert. Plates (Corning lifesciences Costar, #3470) were cultured at 5% CO_2_, 37°C as indicated. Medium was exchanged every 7 days and conditioned media were stored at -80°C. Mosaic photomicrographs were taken from 3D bronchospheres w/o fibroblasts and from GFP expressing primary fibroblasts by bright field and fluorescence microscopy using Axio Observer Inverted microscope/Zeiss^®^ and processed by ZEN microscope navigation Software. Fibroblast proliferation was quantified by mean fluorescence intensity (MFI) which was measured by ImageJ Software as recently described ^12^.

***Treatment of 3D bronchospheres with saracatinib, nintedanib and pirfenidone***

3D bronchospheres were cultivated in a transwell system in matrigel w/wo treatment with vehicle, saracatinib (in a concentration of 600 nM, 210 nM, 75 nM, 25 nM or 8 nM; kindly provided by AstraZeneca)), pirfenidone (1 mM, Santa Cruz Biotechnologie, #sc-203663), nintedanib (1 µM, BIPF 1120, Santa Cruz Biotechnologie, sc-364433) or PBS (control). Medium was exchanged every 7 days.

***Detection of bronchosphere counts and measurement of cell proliferation***

Numbers of bronchospheres per well were counted by bright field microscopy on day 21. Only spheres with a size of 5 µm or larger were counted. Cell proliferation was quantified using the colorimetric MTT assay (Sigma Aldrich, #CT01) on day 21. Briefly, 70 μl of MTT (3-(4,5-dimethylthiazol-2-yl)-2,5-diphenyl tetrazolium bromide) were added into the medium covering the organoids, gently resuspended and then incubated at 37°C for 4h. Then 100 µl isopropanol in 0.04 N HCL were added and the well plate was placed on an orbital shaker (GENEO BioTechProducts GmbH, #1120-221-01) at 150 rpm. Isopropanol dissolves the formazan crystals to a homogenous blue solution. Optical density (OD) was determined at a wavelength of 570 nm using the Infinite 200 PRO Tecan microplate reader (Tecan Group Ltd., INF-FPLEX).

***Calcein AM/ethidium homodimer-1 (“LIVE⁄DEAD^®^”) staining of 3D bronchospheres***

Viability of cells in the 3D bronchospheres was tested by fluorescence microscopy at day 1, 3, 7, 14 and 21 using the Live Death Staining kit. 3D organoids w/wo fibroblasts were incubated with 2 μM Calcein AM (Thermo Fisher, #C1430) and 5 μM Ethidium homodimer-I (EthD-1) (Invitrogen, Thermo Fisher, #E1169) for 45 min at RT in the dark on an orbital shaker at 150 rpm as recently described ^13^. Dye solution was removed and bronchospheres w/wo fibroblasts were washed three times with 300 µl PBS by shaking at 150 rpm for 5 min at RT in the dark on an orbital shaker. Mosaic photomicrographs were taken by fluorescence microscopy using Axio Observer Inverted microscope/Zeiss and ZEN microscope navigation Software. Calcein was detected in the Cy2 channel and Ethidium homodimer in the Cy5 channel.

***Cryopreservation of 3D bronchospheres***

For histology, immunohistology and confocal laser microscopy, 3D organoids were cryopreserved using Tissue Tek O.C.T.^TM^ compound (Hartenstein, #TTEK). The bottom of the transwell insert were excised with a scalpel and embedded in 2 ml Tissue Tek O.C.T.TM compound and placed in a cryomold (Hartenstein, #CMM). Cryomolds containing the 3D organoids were incubated on dry ice for 15 minutes. Cryopreserved bronchospheres were stored at -80°C. Cryosections were prepared using the cryotome Leica CM 1900 (Leica Biosystems Cryostats).

***Measurement of collagen production by Sircol™ Collagen Assay***

Conditioned medium and conditioned matrigel of sphere cultures were harvested after 63 days and stored at -80°C. The conditioned media and matrigels of 3D bronchosphere cultures were used for Sircol assay (Sircol™ Collagen Assay, Biocolor), which was performed as recommended by the manufacturer and recently described ^14^. Briefly, conditioned media and matrigels were digested using the isolation and concentration reagent (polyethylene glycol in a TRIS-HCL buffer pH 7.6) in low binding protein tubes (Thermo Fisher, #90410) to release collagen at 4°C overnight. 1 ml Sircol Dye Reagent was added and incubated on an orbital shaker at 150 rpm for 30 min. During this time period collagen-dye complexes were formed and precipitated out. Afterwards the tubes were centrifuged and washed with ice-cold Acid Salt Wash Reagent (acetic acid, sodium chloride and surfactants) once. Then the collagen dye complexes were dried at RT. Finally, collagen bound dye were dissolved by alkali reagent (0.5M sodium hydroxide) and optical density (OD) of samples were determined using the Infinite 200 PRO Tecan microplate reader at a wavelength of 555 nm.

***Amphiregulin detection by enzyme-linked immunosorbent assay (ELISA)***

Conditioned media of 3D bronchosphere cultures were harvested after 14 days of cell culture and stored at -80°C. Amphiregulin levels of the conditioned media were determined by ELISA following the manufacturer’s instructions (R&D Duosets, #DY262). OD were determined at 450 nm (reference wavelength 540 nm) using a Tecan reader Infinite 200 PRO (Crailsheim, Germany).

***Single cell RNA sequencing of bronchial brushes***

Single cell RNA sequencing (scRNAseq) was performed on bronchial epithelial cells derived from nine patients with IPF and six patients with nonUIP fibrotic ILDs. For baseline characteristics of IPF patients and nonUIP ILD patients see table S1. For scRNAseq experiments, three bronchial brushes were immediately placed in 2 ml of pre-warmed (37°C) BEGM media. Cells were detached from the brush by flicking the brush. Then, airway epithelial cells were pelleted by centrifugation (250×g, 5 min) and disaggregated by resuspension in 0.05% trypsin/EDTA for 5 min at 37°C. Trypsinization was stopped by addition of HEPES buffered saline supplemented with 15% FBS, and the cells were again pelleted at 250×g, 5 min. Cells were frozen in freezing solution (20% DMSO + 80% FBS and BEGM media, 1:1) in liquid nitrogen.

***Sample preparation for single cell RNA sequencing***

Frozen cell suspensions were thawed at 37°C, diluted with 20 ml cold (4°C) PBS (Life Technologies, #14190250) + 10% heat-inactivated FBS (Life Technologies, #10437028), USA), then centrifuged at 300xg at 4°C for 5min. The supernatant was discarded, then the cell pellet was resuspended in PBS + 0.04% BSA (New England Biolabs, # B9000S), and filtered through a 40 μm cell strainer (Fisher Scientific, #50-828-736). For cell concentrations, cells were stained with Trypan blue and counted on the Countess Automated Cell Counter (Thermo Fisher, USA).

***Single cell barcoding, library preparation, and sequencing***

Single cell sequencing was performed using on the 10x Genomics Chromium technology according to the manufacturer’s protocol (Single Cell 3’ Reagent Kits v2, 10x Genomics, USA): Cell suspensions, reverse transcription master mix and partitioning oil were loaded on a single cell “A” chip, then run on the Chromium Controller. mRNA was reversed transcribed to cDNA within the droplets at 53°C for 45 min. cDNA was amplified for 12 cycles in total on a BioRad thermocycler. SpriSelect beads (Beckman Coulter, USA) were used for size selection of cDNA based on a ratio of SpriSelect reagent volume to sample volume of 0.6. For quality control, cDNA was analyzed on Agilent Bioanalyzer high sensitivity DNA chips. cDNA was enzymatically fragmented for 5 min at 32°C, followed by end repair and A-tailing at 65°C for 30 min. A double-sided size selection of the cDNA was performed using SpriSelect beads. After adaptor ligation, cDNA was amplified using a sample-specific index oligo as primer, followed by a last double-sided size selection using SpriSelect beads. Final cDNA libraries were again analyzed on an Agilent Bioanalyzer high sensitivity DNA chip. Libraries were sequenced on a HiSeq 4000 Illumina platform aiming for 150 million reads per library.

***Data processing***

Base calls were converted to fastq reads using the wrapper ‘mkfastq’ of the software Cell Ranger (10x Genomics, USA). Template switch oligo sequence (AAGCAGTGGTATCAACGCAGAGTACATGGG) and poly(A) sequence contaminations on Read2 were trimmed with cutadapt (v2.3) ^15^. If Read2 sequences were shorter than 30 bp after trimming, read pairs were discarded. zUMIs pipeline (v2.0)^16^ was used for subsequent processing of reads. Paired reads were filtered if either the cell barcode or unique molecular identifier (UMI) sequence had more than 1 bp with a phred < 20. Read2 sequences were mapped to the human genome reference GRCh38 release 91 from ensemble ^17^ using STAR (v2.6.0c) ^18^. Reads were collapsed on a UMI level, reads that aligned to both exonic and intronic sequences were retained as both separate and combined gene expression assays. To delineated cell barcodes representative of quality cells from barcodes of apoptotic cells or background RNA, the following thresholds had to be passed: having at least 17.5% of transcripts arising from intronic, i.e. unspliced reads, indicative of nascent mRNA; profiling of more than 3000 UMIs per cell barcode; and less than 20% of their transcriptome being of mitochondrial origin. Gene identities were output in ensemble gene ID format. To improve the interpretability of the gene identities, ensemble gene IDs were converted to HGNC format only if an exact one-to-one translation was available using the R package BioMart ^19^.

***Computational Analysis***

Raw UMI counts were normalized with a scale factor of 10,000 UMIs per cell, and then natural log transformed using a pseudocount of 1. Louvain cluster analysis ^20^ was performed to identify cell types using the R package Seurat (version 3) ^21^. Multiplet cell populations, identified as having a transcriptomic gene expression profile that resembled the resulting combination of 2 disparate cell type signatures that already existed in the dataset, and low UMI cell populations were not included in downstream analyses ^22^. Highly variable genes in the dataset were identified as the top 1000 genes ranked by dispersion (scaled variance/mean) across 20 bins of the expression distribution. The expression values for these genes were then adjusted for differences in total UMI and the fraction of mitochondrial reads across cells during z-normalization with a maximum absolute z-score of 10. For linear dimension reduction, these scaled gene expression values were subject to principle component analysis (PCA). A shared nearest neighbor network was created based on Euclidean distances between cells in multidimensional PC space and a fixed number of neighbors per cell, which was used to generate a 2-dimensional Uniform Manifold Approximation and Projection UMAP ^23^ for visualization. UMAP embeddings of the full dataset are depicted in figure S4. The dataset was then subsetted to all epithelial populations, re-embedded in UMAP space and colored by cell type and disease status (Figure 3A,B). To account for subject variability, unity normalized gene expression of canonical marker genes of the epithelial cell types, averaged per subject, grouped by cell type, was visualized in a heatmap (Figure 3C). Gene expression between groups of cells was compared using a Wilcoxon Rank Sum test with Bonferroni correction of p-values for multiple testing. Significantly differentially expressed, ugregulated genes in IPF basal cells compared to control basal cells were subjected to a pathway analysis using the software enrichr and the human KEGG 2019 pathway database ^24, 25^.

***Connectivity Map (CMap) analysis to identify pertubagen classes reversing IPF-associated gene expression profiles of ABCs***

Broad Institute's CLUE platform (<https://clue.io>) was used to identify a potential molecular mechanism of action which could reverse A) the deviating gene expression profile of ABCs in IPF as identified by scSeq in this study and B) the ABC gene expression signature associated with mortality in IPF by bulk RNA sequencing as recently described by us ^26^. CLUE is based on the Connectivity Map data set ^27^ that analyses cellular gene expression responses to multiple perturbations. As input, we used A) the top 150 (the maximum number of inputs in CLUE) genes differentially higher expressed in IPF vs nonUIP ILD Controls in the ABC population of our scSeq dataset ordered by FDR and B) the top 150 genes associated with mortality from Prasse et al.^26^ ordered by FDR. Furthermore, all input genes had to be comprised in the gene space of CLUE (<https://clue.io/command?q=/gene-space%20lm>). The analysis was run in CLUE’s data version 1.1.1.2 and the software version 1.1.1.41. The summary connectivity score, ranging from +100 (pertubagen classes with transcriptional effects similar to the input gene signature) to -100 (perturbagen classes with the opposite effect to the input gene signature, i.e. potential treatments) of all pertubagen classes was extracted, and visualized as bar plots of the summary connectivity scores split by the origin of the input gene profiles.

***Experimental protocol for the humanized mouse model for IPF***

All animal models were performed multiple times as indicated and data were jointly analyzed. All animals treated were included in the analysis. Interventions were not blinded, but analysis of animal samples was blinded, and, in each experiment, respective controls were included. *Rag2^-/-^*, *NRG* and *NSG* mice received a dose of 1.2 mg/kg bleomycin intratracheally at day 0 during light halothane-induced anesthesia. Three days later 0.3 x 10^5^ (*NRG* and *NSG* 0.2 x 10^5^) human ABCs from patients with IPF, nonUIP ILD or healthy volunteers were intratracheally injected during light halothane-induced anesthesia. In some experiments lentiviral transduced human ABCs were used (see below). In addition, in some experiments mice were treated oropharyngeally with 10 mg/kg of saracatinib (kindly provided by Leslie Cousens, AstraZeneca) daily versus vehicle (200 µl PBS) daily. For these experiments we used pairs of *NRG* mice in which the same human IPF-ABC line was injected. One mouse of the pair was treated with saracatinib and the other with vehicle (PBS).

Unless stated otherwise, lungs were harvested at day 21. For histological and immunohistological analyses, the trachea was cannulated, and lungs were insufflated with 4% paraformaldehyde in PBS at a pressure of 25 cm H_2_O, followed by removal of the heart. Inflated lungs were incubated in 4% formaldehyde solution overnight at 4°C and after this step 4% formaldehyde solution were replaced by 70% ethanol. Lungs were stored in 70% ethanol at RT until paraffin embedding. For hydroxyproline measurements left and right lungs were weight and homogenized in distilled water on a ULTRA-TURRAX^®^ (VWR, #IKAA3725001) and stored at -80°C until used.

***Hydroxyproline and total protein assay***

Murine lung hydroxyproline was determined by the hydroxyproline colorimetric assay kit from Biocat (hydroxyproline Kit; Biocat GmbH, #K55-100) following manufacturer’s instruction, as previously described ^28^. Lung homogenates (100 µl) were mixed with 100 µl 12 N HCl, and the samples were incubated (hydrolyzed) at 120°C in a pressure-tight, teflon-capped vial (Merck, #Z115096) for 3 hours. Afterwards 100 µl of the chloramine T reagent were added to each sample and incubated at RT for 5 min. Then 100 μl of DMAB reagent were added to each well and incubated for 90 min at 60°C. OD of each sample was determined using the Infinite 200 PRO Tecan microplate reader at a wavelength of 560 nm. Total protein levels were analyzed by the total protein assay kit of Quickzyme (Netherlands, #QZBTOTPROT1) according to the manufacturer`s description. Briefly: hydrolyzed lung homogenate samples were diluted 1:2 in 6 M HCL. Then 15 µl of the standard solution (3000 µg/ml high standard, 0.047-3.00 mg/ml) and 15 µl of the samples were added in a 96 well microplate and 120 µl of the assay buffer were added, then mixed on a shaker at 150 rpm for 10 min at RT. Afterwards 15µl of the color working reagent solution were added to each well. The microplate was covered with an adhesive film and incubated at 85°C for 60 min in an incubator. OD was determined at 570 nm using the Infinite 200 PRO Tecan microplate reader. Hydroxyproline data are expressed as hydroxyproline [µg/ml]/ total protein [mg/ml].

***Testing engraftment of human ABCs in mouse lungs by bioluminescence imaging***

For *in-vivo* imaging of FLuc transduced human ABCs in NRG mice, 150 mg/kg XenoLight D-Luciferin-K+ Salt Bioluminescent Substrate (PerkinElmer, #122799) was injected subcutaneously on day 7, 14, and 21. Mice were anesthetized (1.5% to 2.5% isoflurane) and bioluminescence was measured by an IVIS Lumina II (PerkinElmer, USA). Data were analyzed using LivingImage 4.5 (PerkinElmer, USA).

***Lentiviral vector for firefly-luciferase and enhanced green fluorescent protein (eGFP) overexpression***

The lentiviral vector pRRL.PPT.CBX3-SFFV.FLuc.T2A.eGFP.P2A.Neo.pre (GFP vector) was generated by cloning of the cDNAs for firefly-luciferase (FLuc), enhanced green fluorescent protein (eGFP) and neomycin (Neo) into a lentiviral vector backbone (kindly provided by Dr. Axel Schambach).

***Lentiviral vector for c-src overexpression and knockout***

The vector pRRL.PPT.CBX3-SFFV.C-SRC.E2AFLuc.T2A.eGFP.P2A.Neo.pre (c-SRC OE vector) for overexpression of c-src was cloned by insertion of a codon-optimized c-src cDNA in front of FLuc into the pRRL.PPT.CBX3-SFFV.FLuc.T2A.eGFP.P2A.Neo.pre vector.

Knockout of c-src was achieved using an all-in-one CRISPR-Cas9 vector pRRL.PPT.hU6.C-SRC-sgRNA.SFFV.spCas9-2xNLS.T2A.dTomato.pre (c-SRC KO vector) (kindly provided by Dr. Axel Schambach). The sgRNA target sequence for c-src was cloned after phosphorylation and annealing of the oligonucleotides 5’-CACCGAGCGCCGTGCACGTTCTCGG and 5’-AAACCCGAGAACGTGCACGGCGCTC via two BsmBI sites into the all-in-one CRISPR-Cas9 vector.

***Virus production and titration***

Lentiviral vector supernatants were produced by transfection of 10 µg lentiviral vector, 12 µg gag-pol, 6 µg rev and 2 µg VSVg packaging plasmids into 293T cells (ATCC, LGC Standards, Wesel, Germany) in 10 cm plates. All plasmids were mixed and transfected using calcium-phosphate in the presence of 25 µM chloroquine (Sigma-Aldrich, Seelze, Germany, #C6628). Media exchange was performed 6 hours and viral supernatants were harvested 30 and 54 hours after transfection. Pooled viral supernatants were filtered through 0.22 μm filters (Millex-GP, Millipore, #SLGP033RS) and 100-fold concentrated using an ultracentrifugation step for 2 hours at 25,000 rpm and 4°C. The infectious titer was determined by applying serial dilutions of viral supernatants onto 10^5^ HT1080 cells (DSMZ, Braunschweig, Germany) in the presence of 4 µg/ml protamine sulfate (Sigma-Aldrich, #P3369).

***Transductions***

For transduction of ABCs and fibroblasts, cells were plated with 30 % confluency in T25 flasks 14 days before transduction. ABCs were transduced with c-SRC OE, c-SRC KO, eGFP lentiviral vector supernatants and in addition fibroblasts with eGFP lentiviral vector supernatants. ABCs were cultivated in BEGM media and fibroblasts in DMEM media. Transduction was performed in the presence of 4 µg/ml protamine sulfate (Sigma-Aldrich, #P3369) using a multiplicity of infection (MOI) of 0.25 - 0.5. Media exchange was performed 6-8 hours after transduction with BEGM and DMEM media as previously described ^29^. Two days later, transduced cells were selected by applying 1 mg/ml G418 (geneticin, selective antibiotic, Lonza, #15-394N) in BEGM media and DMEM media. Selection media were changed every three days. Luciferase activity of cells was tested using a luminometer prior to use.

***Histology and immunohistochemistry***

Immunohistochemistry (IHC) of formalin fixed lung tissues from 10 patients with IPF (explants), and 3 healthy lung donors (transplants) was performed. In addition, formalin-fixed mouse lung tissues of the described humanized IPF mouse model and cryopreserved 3D organoids were used for immunohistochemistry. Tissue sections (3 µm) of formalin-fixed human and murine lung tissues embedded in paraffin blocks were deparaffinized in xylene and rehydrated with a descending alcohol row: 3 times 100% Xylene (Merck, #108633), 2 times 100% ethanol (Merck, #107017), once 90% ethanol, once 70% ethanol and distilled water each for 3 min. Heat induced antigen retrieval was performed at 110°C for 2 min in citrate buffer (pH 6.0) using a pressure cooker. Prior to staining, slides were rinsed for 5 min with Tris-NaCl buffer and were blocked for 20 min with normal goat serum 1:20 (Vector, #S1000). Incubations with primary antibody were conducted 30 min at RT. All primary antibodies and dilutions are listed below. Afterwards, slides were rinsed with Tris-NaCl buffer. Secondary biotinylated antibodies were incubated for 30 min at RT. Afterwards slides were rinsed with Tris NaCl buffer. Activation was done using alkaline phosphatase Strept AP (1:800 dilution) (Vector, #SA-5100) for 30 min and slides were rinsed with Tris-NaCl buffer. Visualization was performed by DAKO REAL Chromogen Red (Dako Real Kit) (Dako/Agilent Technologies, #K500311-2) (incubation time: 20 min). Afterwards, slides were rinsed with distilled water. Counterstain with mayer`s hemalum solution (Merck, #109249) 1:10 for 90 sec. Slides were rinsed shortly with distilled water and for 90 sec with Shandon™ Bluing Reagent (ThermoScientific, #6769001). Before mounting, slides were rinsed 1 time with distilled water followed by 3 times with 90% ethanol, 6 times with 100% ethanol and 6 times with xylene. Finally, slides were coverslipped with Eukitt Quick-hardening mounting medium (Merck, #03989). 3D Organoids were stained directly with three primary antibodies against CK5/6, p63 and αvβ6 integrin. The IHC staining takes place immediately as described above with the exception that the endogenous peroxidase was deactivated by H_2_O_2_ (Merck, #107209) for the second and third primary antibody. Visualization was performed with three different Chromogens DAKO REAL Chromogen Red, DAKO FLEX HRP turquoise Chromogen and EnVision™ FLEX DAB+ brown Chromogen DAKO with Peroxidase coupled secondary antibody (DAKO EnVision FLEX/HRP). All slides were digitalized using Mirax Scan 150 BF/FL (Zeiss, Germany). Details and antibodies of the methods are provided in the table below.

**Primary antibodies and methods used for IHC**

**2**

| **Marker** | **Clone** | **Vendor** | **Procedure** | **Dilution/ Conc.** | **Incubation** | **Detection** |
| --- | --- | --- | --- | --- | --- | --- |
| C-src | polyclonal anti-src rabbit antibody | Abcam, #ab47405 | Manual | 1:50 | 30 min, RT | AP, (DAKO REAL Streptavidin Alkaline Phosphatase) |
| CK5/6 | monoclonal mouse anti-human CK5/6 (DAKO clone D5/16 B4) | Dako  Omnis, #GA78061-2 | Manual | Ready to use | 30 min, RT | AP, (DAKO REAL Streptavidin Alkaline Phosphatase) |
| P63 | P63 polyclonal rabbit anti-human antibody | Calbiochem, #PC373 | Manual | 1:1000 | 30 min, RT | HRP, Peroxidase coupled secondary antibody (DAKO EnVision FLEX/HRP) |
| anti-human anti-αvβ6 | MAb6.3G9, primary monoclonal rat antibody | kindley provided by Shelia Violette Biogen Idec | Manual | 1:100 | 30 min, RT | HRP, Peroxidase coupled secondary antibody (DAKO EnVision FLEX/HRP) |
| negative control | Negative Control Mouse IgG1 antibody | Dako Omnis,  #GA75066-2 | Manual | 1:20 | overnight,4°C | AP, (DAKO REAL Streptavidin Alkaline Phosphatase) |
| negative control | FLEX Universal Negative Control, Rabbit | Dako Omnis,  #IS60061-2 | Manual | 1:20 | overnight,4°C | AP, (DAKO REAL Streptavidin Alkaline Phosphatase) |
| negative control | Negative Control Mouse IgG1 antibody | Dako Omnis,  #GA75066-2 | Manual | 1:20 | overnight,4°C | AP, (DAKO REAL Streptavidin Alkaline Phosphatase) |

**Secondary antibodies** **and methods used for IHC**

| **Clone** | **Vendor** | **Procedure** | **Dilution/ Conc.** | **Incubation** |
| --- | --- | --- | --- | --- |
| Goat IgG anti-Mouse IgG (H+L)-Biotin | Dianova, #115-065-003 | Manual | 1:800 | 30 min RT |
| Goat IgG anti-Rabbit IgG (H+L)-Biotin | Dianova, #111-065-003 | Manual | 1:800 | 30 min RT |

***Masson’s trichrome staining and fibrosis scoring***

Masson trichrome stains were performed from formalin-fixed paraffin-embedded murine lung tissues according to a standard protocol as recently described ^13^. In brief, deparaffinization of paraffin sections mounted on slides was performed in a cuvette with a descending alcohol row: 3 times 100% xylene, 2 times 100% ethanol, once 90% ethanol, once 70% ethanol for 3min and then incubated in distilled water for 10 min. Then slides were incubated in bouin's solution (150 ml picric acid solution 1.3% (Sigma-Aldrich, #P6744), 50 ml formaldehyde 37% (Carl Roth, #CP10.2) and 10 ml ice acetic acid (Merck, #1.0056) for 15 min at 56°C. Afterwards slides were cooled at RT and rinsed under tap water. Then slides were incubated in Weigert's solution A and B (Weigert's iron hematoxylin kit; Merck, #1.15973.1) for 10 min. Slides were rinsed under tap water and distilled water and dyed for 5 min in Biebrich scarlet-acid fuchsin solution (Sigma-Aldrich, #HT151). After another washing step slides were incubated for 10 min in molybdatophosphoric acid solution (Sigma Aldrich, #1.00532). Afterwards slides were directly transferred to aniline blue Chroma (Sigma Aldrich, #B8563). Then, slides were rinsed with distilled water, acetic acid and dehydrated with a descending alcohol row: 70% ethanol 2 min, 96% ethanol 2 min, 100 % ethanol 2 min and 100% xylene 10 min. At the end slides were mounted with Roti^®^-Histokitt II (Carl Roth, #160.1). Left lung sections were imaged using Mirax Scan 150 BF/FL (Zeiss, Germany). Ashcroft score was calculated as previously described by Ashcroft and us ^30, 31^. The scoring scale was as follows: 0 = no abnormalities, 1 = slight thickening of alveolar membranes, 2 = small areas of fibrosis (<10%), 3 = 10–20% fibrotic area, 4 = 20–40% fibrotic area, 5 = 40–60% fibrosis, 6 = 60–80% fibrosis, 7 = >80% fibrosis, and 8 = complete fibrosis.

***Hematoxylin-eosin (H&E) staining***

H&E staining was performed according to a standard protocol and as recently described.^32^ Sections were incubated in Mayer's hemalum solution (Sigma Aldrich, #109249) for 2 min followed by rinse in PBS for 3 times and then stained with Eosin Y (yellowish) (C.I. 45380) (Sigma Aldrich, #115935) in combination with Phloxin B (C.I. 45410) (Sigma Aldrich, #115926) for 20 sec. H&E staining were carried out in the H&E slide stainer Leica Biosystems #STS5020-005030. Slides were digitalized using Mirax Scan 150 BF/FL (Zeiss, Germany).

***Confocal laser scanning microscopy (CLSM)***

CLSM of representative cryopreserved 3D organoids w/wo IPF fibroblasts, representative human lung tissue sections and representative murine lung tissue sections was performed. 3µm thick sections of paraffin-embedded blocks were deparaffinized with a descending alcohol row as described. Heat induced antigen retrieval was performed at 110°C for 2 min in citrate buffer (pH 6.0) using a pressure cooker. At first, staining slides were washed for 5 min with Tris-NaCl buffer and then blocked for 20 min with 4% normal donkey serum (Dianova, #017-000-121). Of cryopreserved 3D organoid cultures 20 µm thick sections were obtained by Leica CM 1900 (Leica Biosystems, Germany) and directly fixed with acetone (Carl Roth, #5025.1) for 15 min at -20°C and then dried at RT. Slides were stored at -20°C until re-use. Prior to immunofluorescence staining slides were rehydrated for 5 min with PBS and then blocked for 15 min with 4% normal donkey serum (Dianova, #017-000-121).

For immunofluorescence analysis, sections were stained with primary and secondary antibodies as listed below. The primary antibodies were incubated at 4°C overnight in a humidified chamber. On the following day, slides were washed 3 times with 2 ml PBS and the secondary antibodies were incubated for 2 hours at RT. Nuclei were stained by DAPI (Sigma Aldrich, #D9542) as recommended by the manufacturer. Finally slides were washed 3 times with 2 ml PBS and once with 2 ml distilled water and mounted with prolong diamond antifade (Thermo Fisher Scientific, #P36970). Pictures were taken at a laser scanning microscope Olympus FluoView 1000 and with a LSM 510 Meta/Zeiss laser scanning microscope (CLSM). Images were analyzed and processed with software Imaris 7.6 Bitplane Scientific (Oxford Instruments company/USA).

**Primary and secondary antibodies used for immunofluorescence staining**

**2**

| **Sections** | **Marker** | | **Clone** | | **Vendor** | | **Dilution** | | **Detection Ab** | | **Vendor** | |
| --- | --- | --- | --- | --- | --- | --- | --- | --- | --- | --- | --- | --- |
| Human lung tissue | | c-src | | polyclonal anti-src rabbit antibody | | Abcam,  #ab47405 | | 1:400 | | Donkey IgG anti-Rabbit IgG (H+L)-Cy3 | | Dianova,  #715-547-003 |
| Human lung tissue | | CK5/6 | | FLEX Monoclonal Mouse Anti-Human Cytokeratin 5/6 Clone D5/16 B4 | | Dako Omnis,  #GA78061-2 | | Ready-to- Use | | Donkey Fab anti-Mouse IgG (H+L)-Alexa Fluor 488, Cy2 | | Dianova,  #715-547-003 |
| NSG-Venus Mice lung tissue | | CK8 | | Rabbit Anti-Cytokeratin 8 antibody [EP1628Y] | | Abcam, #ab53280 | | 1:100 | | Donkey IgG anti-Rabbit IgG (H+L)-Cy3 | | Dianova, #711-165-152 |
| NSG-Venus Mice lung tissue | | CK5/6 | | FLEX Monoclonal Mouse Anti-Human Cytokeratin 5/6 Clone D5/16 B4 | | Dako Omnis, #GA78061-2 | | Ready-to-Use | | Donkey IgG anti-Mouse IgG (H+L)-Cy5 | | Dianova, #715-175-151 |
| NRG mice lung tissue | | AQP5 | | Goat AQP5 Antibody (G-19) | | Santa Cruz, #sc-9890 | | 1:25 | | Donkey IgG anti-Goat IgG (H+L)-Cy3 | | Dianova, #705-165-147 |
| NRG mice lung tissue | | eGFP | | GFP (D5.1) XP® Rabbit mAb | | Cell Signalling, #2956 | | 1:75 | | Donkey Fab anti-Rabbit IgG (H+L)-Alexa Fluor 488 | | Dianova, #711-547-003 |
| NRG mice lung tissue | | CK5/6 | | FLEX Monoclonal Mouse Anti-Human Cytokeratin 5/6 Clone D5/16 B4 | | Dako Omnis, #GA78061-2 | | Ready-to-Use | | Donkey Fab anti-Mouse IgG (H+L)-Alexa Fluor 488 | | Dianova, #715-547-003 |
| Cryo Slide/ 3D Organoids w/o IPF fibroblasts | | CK5/6 | | FLEX Monoclonal Mouse anti-human cytokeratin 5/6 Clone D5/16 B4 | | Dako Omnis, #GA78061-2 | | Ready-to-Use | | Alexa Fluor® 488- conjugated AffiniPure Fab fragment donkey anti mouse IgG | | Dianova, #715-547-003 |
| Cryo Slide/ 3D Organoids  w/o IPF fibroblasts | | Anti-alpha smooth  muscle Actin | | Rabbit monoclonal | | Abcam,#  ab124964 | | 1:400 | | Cy™5 AffiniPure DonkeyAnti-RabbitIgG (H+L) | | Jackson immuno research, #711-175-152 |
| Cryo Slide/ 3D Organoids w/o IPF fibroblasts | | Vimentin | | goat | | Merck,  #V4630 | | 1:20 | | Donkey IgG anti-Goat IgG (H+L)-Cy3 | | Dianova, #705-165-147 |
|  | | Isotype control | | Rabbit IgG, polyclonal – Isotype Control | | Abcam, #ab37415 | | 1:400 | | Cy™5 AffiniPure DonkeyAnti-RabbitIgG (H+L) | | Jackson immuno research, #711-175-152 |
|  | | Negative control | | FLEX Universal Negative Control, Mouse | | Dako Omnis, #GA75066-2 | | Ready-to- Use | | Donkey Fab anti-Mouse IgG (H+L)-Alexa Fluor 488, Cy2 | | Dianova, #715-547-003 |
|  | | Isotype Control | | Goat IgG Isotype Control | | Thermo Fisher Scientific, #31245 | | 1:25 | | Donkey IgG anti-Goat IgG (H+L)-Cy3 | | Dianova, #705-165-147 |

***Protein extraction and western blot analysis***

Proteins from tissues or cultured cells were extracted/lysed in ice-cold RIPA buffer containing 1M NaCl (Merck, #106404), 1% Nonidet P40 (Merck, #11754599001), 0,5% sodium deoxycholate (Merck, #D6750), 0,1% SDS (Merck, #L3771), 50 mM Tris (Carl Roth, #4855.2) pH 7.4 in ddH_2_O with protease/phosphatase inhibitor (Cell Signaling, #5872) and incubated on ice for 20 min. Protein concentrations were determined by BCA Kit (Thermo Fisher Scientific, #A53225) as recommended by the manufacturer. Samples were incubated 5 min in Laemmli-sample buffer (BioRad, #1610747) containing 10% β-mercaptoethanol (Merck, #M6250) at 95°C. Samples were separated using 12% SDS-PAGE gels (BioRad, #671044) and proteins transferred to polyvinylidene difluoride membranes by [Trans-Blot^®^ Turbo™ transfer system](https://www.google.com/url?sa=t&rct=j&q=&esrc=s&source=web&cd=1&cad=rja&uact=8&ved=2ahUKEwifie2Mo6LiAhUBy6QKHaGoC5UQFjAAegQIABAB&url=http%3A%2F%2Fwww.bio-rad.com%2Fde-de%2Fproduct%2Ftrans-blot-turbo-transfer-system%3FID%3DLGOQBW15&usg=AOvVaw1TAYr-GoeZgyKUaYW93m17) (BioRad/Germany). After blocking with 5% non-fat dry milk in TBS-T buffer (25 mM Tris-HCl, 150 mM NaCl, 0.1% Tween 20 (Merck, #P1379), pH 7.5), the membrane was incubated overnight at 4°C with one of the following primary antibodies listed in the table below. GAPDH and α-Tubulin were used for all blots as endogenous control. All primary antibodies were detected using secondary antibody goat anti rabbit (H+L)-HRP conjugate (1:3000) (BioRad, #1706515) and secondary goat anti mouse (H+L)-HRP conjugate (1:3000) (BioRad, #1706516). Enhanced chemiluminescence on immunoblot gels (ClarityTM Western ELC Substrate (BioRad, #1705060) was used for detection with ChemiDoc TM MD Imaging System (BioRad/Germany). P-EGF, tot- EGF, α-SMA, Collagen I, P-p44/42 and tot-p44/42 was detected in lysates of healthy donor fibroblasts (Fib). Fib were seeded into a 12-well plate, grown until 80–90% confluence, incubated with serum-free medium overnight, and then exposed to 14 days old conditioned medium of low and big 3D organoids for either 20 min or 48 hours. Afterwards, the cells were directly lysed in 5×Laemmli buffer containing 10% β-mercaptoethanol and subjected to western blotting. Western Blot was performed as described above with the only exception that a 7.5% and 10% SDS polyacrylamide gel were used.

**Primary antibodies used for Western Blot**

| **Marker** | **Clone** | **Vendor** | **Dilution/ Conc.** |
| --- | --- | --- | --- |
| c-src | polyclonal anti-src rabbit antibody | Abcam,  #ab47405 | 1:500 |
| Type I Collagen | polyclonal goat anti-Type I Collagen | Southern Biotec,  #SAB-1310-01 | 1:1000 |
| α-SMA | monoclonal mouse anti-α-SMA, ASM-1 | Merck Millipore,  #CBL171 | 1:2000 |
| Fibronectin,cFn | monoclonal mouse anti-FN, DH1 | Enzo Life Sciences,  #BML-FG6010 | 1:500 |
| p44/42 (Thr202/Tyr204) | mouse anti-phospho-p44/42 (Thr202/Tyr204) | Cell Signaling,  #9106 | 1:500 |
| phospho-EGFR (Tyr845) | rabbit phospho-EGFR (Tyr845) | Zytomed,  #205-0235 | 1:500 |
| total EGFR | rabbit anti-total EGFR | Biomol | 1:500 |
| GAPDH (6C5) | mouse monoclonal antibody raised against GAPDH | Santa Cruz,  #sc-32233 | 1:200 |
| α-Tubulin | Rabbit anti- α-tubulin antibody | Cell Signaling,  #2125 | 1:1000 |

**Secondary antibodies** **used for Western Blot**

| **Clone** | **Vendor** | **Dilution/ Conc.** |
| --- | --- | --- |
| goat anti rabbit (H+L)-HRP conjugate | BioRad, #1706515 | 1:3000 |
| secondary goat anti mouse (H+L)-HRP conjugate | BioRad, #1706516 | 1:3000 |

**Statistical Analysis**

Statistical analysis was performed with GraphPad Prism 8.0 (GraphPad Software Inc). Comparison between groups was done as indicated. In all cases a p-value less than 0.05 was considered significant. Statistical analysis of the single cell data set incl. the software used is described in the methods section, as it is part of the workflow.

**Data and Code Availability**

The single cell sequencing data of human bronchial Brush cells in IPF and disease controls will be uploaded to the GEO database.

**Supplementary Tables**

#### Table S1. Baseline Characteristics of IPF Patients and nonUIP ILD patients of the single cell sequencing cohort

| **Table S1. Baseline Characteristics of IPF and nonUIP ILD Patients*** | | |
| --- | --- | --- |
| **Characteristic** | **IPF**  N=9 | **NonUIP ILD**  N=6 |
| Age -yr | 71 ± 7 | 64 ± 11 |
| Male/ female sex | 7/2 | 6/0 |
| FVC percent predicted value | 71 ± 9 | 69 ± 27 |
| DLCO percent predicted | 54 ± 13 | 53 ± 20 |
| Smoking Status |  |  |
| Never smoked -% | 3 | 3 |
| Former smoker-% | 6 | 3 |
| Current smoker-% | 0 | 0 |

* Plus–minus values are means ±SD. FVC denotes for forced vital capacity. DLCO denotes for diffusion capacity. NonUIP ILD denotes for patients with a fibrotic interstitial lung disease (ILD) who presented with a computer tomography (CT) scan that is not consistent with a definite Usual Interstitial Pneumonia (UIP) pattern. IPF denotes for idiopathic pulmonary fibrosis.

### Supplementary Figures

**Figure S1**

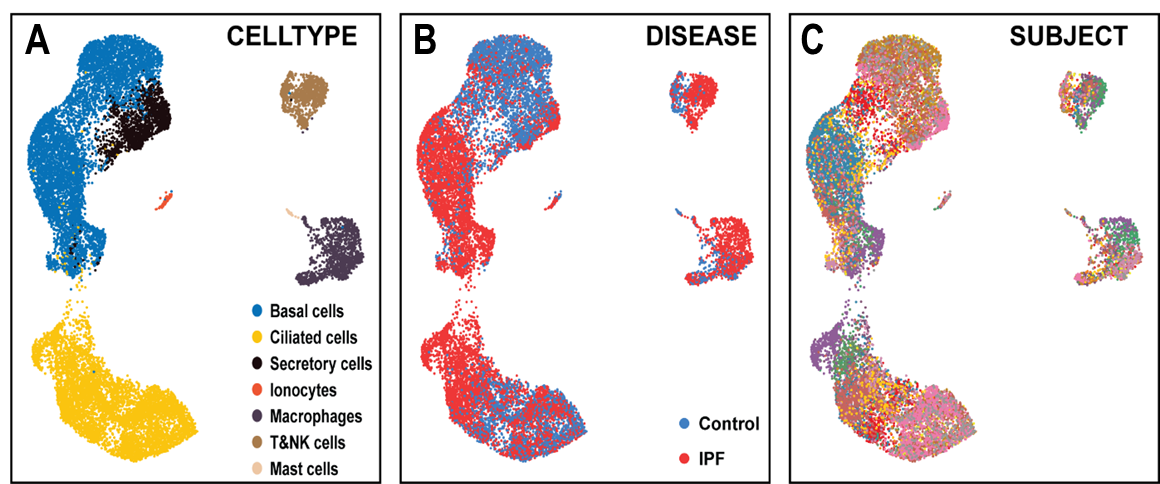

***Figure S1.*** *Uniform Manifold Approximation and Projection (UMAP) of the full dataset of 17,339 single cell transcriptomes visualizes the seven major discrete cell types detected.* ***(A)*** *UMAPs colored by cell types.* ***(B)*** *UMAPs colored by disease states.* ***(C)*** *UMAPs colored by subjects.*

#### Figure S2

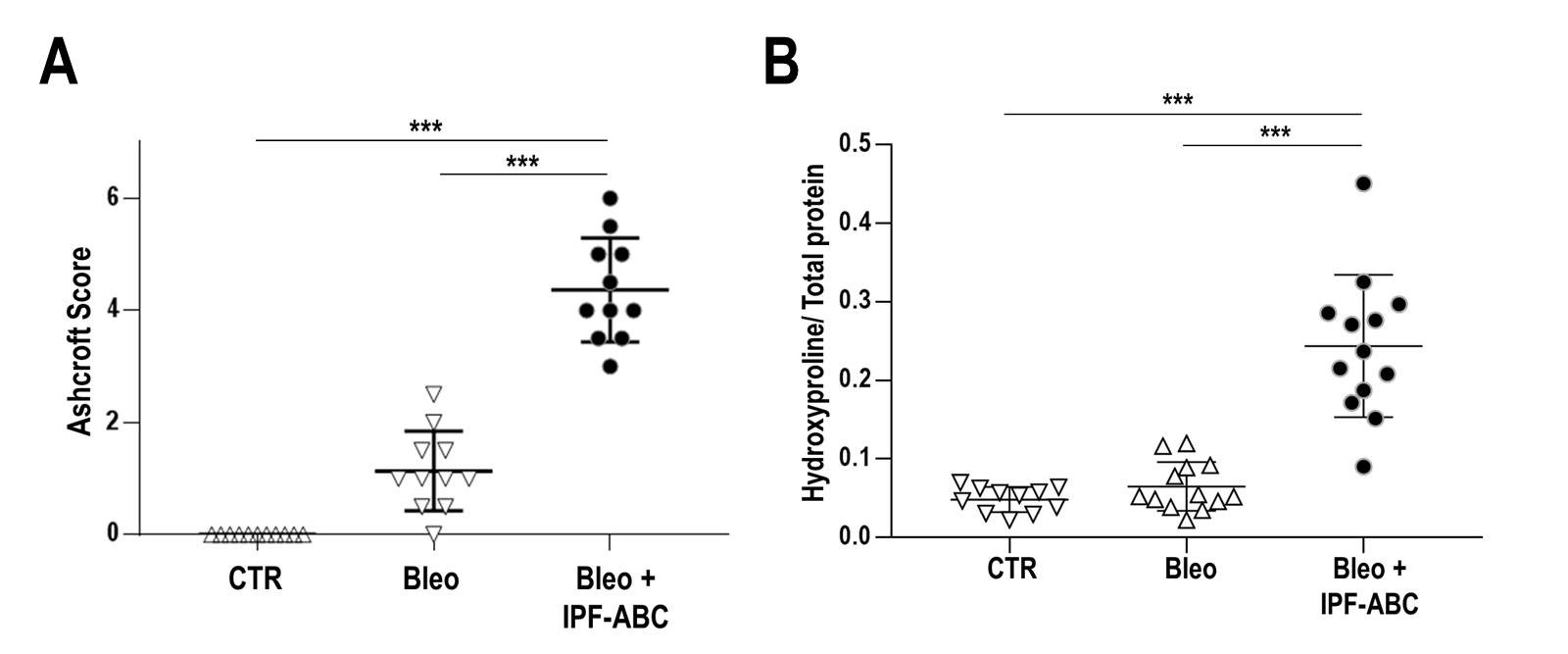

***Figure S2. Airway basal cells of IPF patients (IPF-ABC) augment fibrosis in bleomycin challenged NRG mice.*** *NRG mice received either PBS (CTR) or bleomycin (Bleo) intratracheally and some mice (Bleo + IPF-ABC) additionally three days later 200.000 IPF-ABCs intratracheally. Lungs of mice were harvested for either histopathological scoring (Ashcroft, (****A****) or hydroxyproline measurements (****B****). CTR denotes for control mice; Bleo denotes for bleomycin; IPF-ABC denotes for airway basal cells derived from IPF patients. ***P<0.001, ANOVA with Tukey correction for multiple testing (A,B).*

#### Figure S3

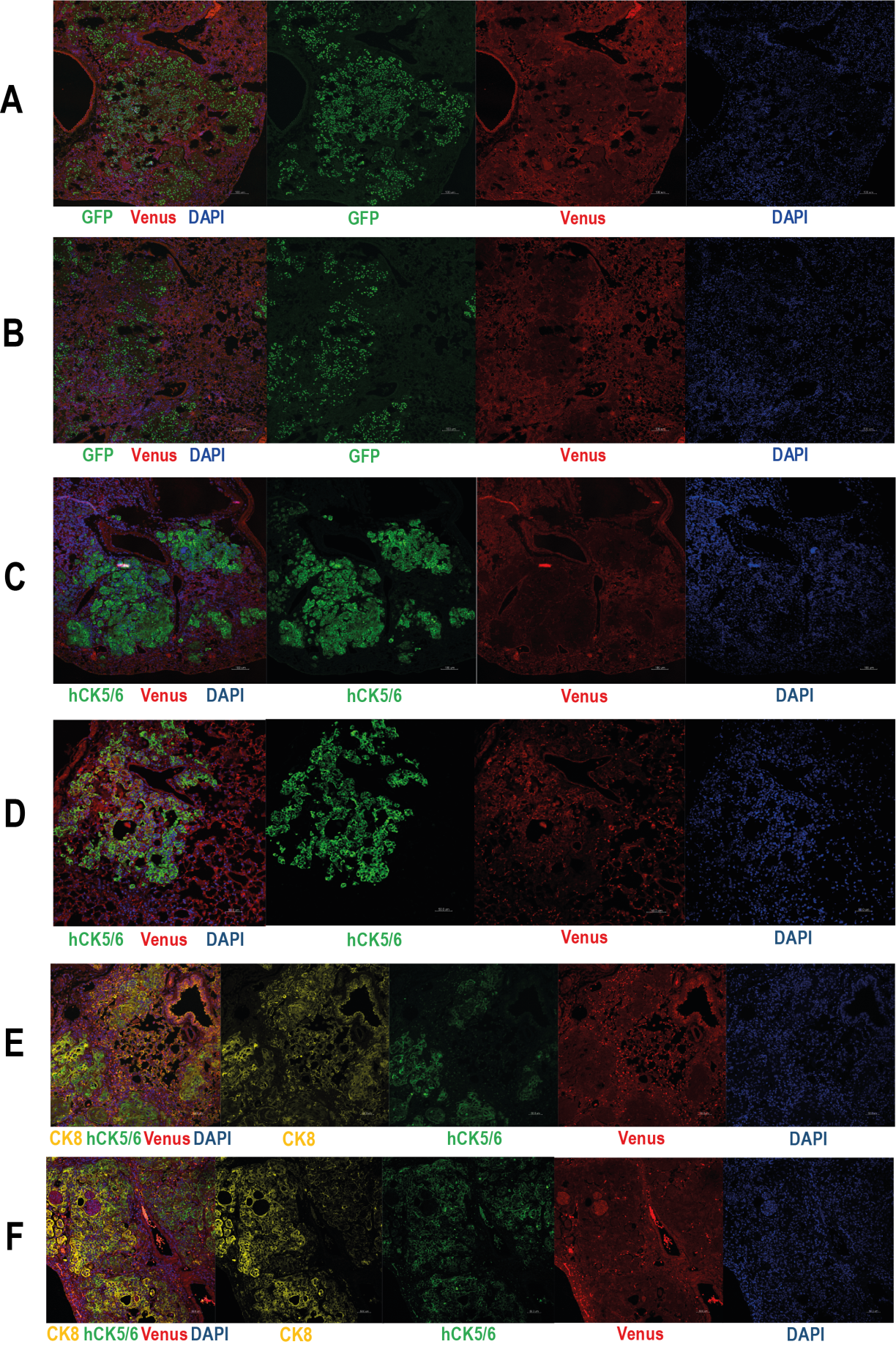

***Figure S3.*** ***Overlay and single original registrations obtained by confocal laser microscopy which are depicted in Figure 2I.*** *NSG mice received bleomycin i.t. at day 0 and three days later human IPF-ABCs which were transduced with a lentiviral vector encoding for eGFP. Lungs were harvested at day 21. (****A-B****) Shown are eGFP staining in green, Venus expression in red, and nuclei staining by DAPI in blue.* ***(C-D)*** *Shown are human cytokeratin 5/6 (hCK5/6) staining in green, Venus expression in red and nuclei staining by DAPI in blue* ***(E-F)*** *Shown are human cytokeratin 5/6 (hCK5/6) staining in green, cytokeratin 8 (CK8) staining in yellow, Venus expression in red and nuclei staining by DAPI in blue. Scale bars, 50µm.*

**Figure S4**

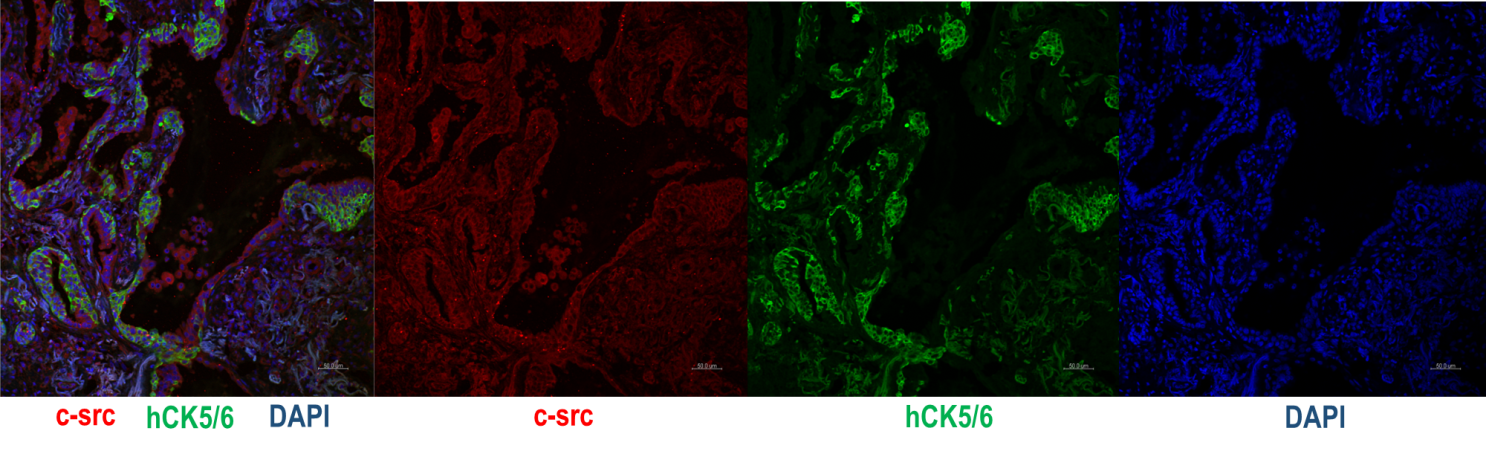

***Figure S4. C-SRC is highly expressed in human lung tissues derived from patients with IPF.*** *Overlay and single original registrations of human IPF lung tissue obtained by confocal laser microscopy which are depicted in Figure 4B. CK5/6 staining signal in green, c-src staining in red and nuclei staining by DAPI in blue.*

#### Figure S5

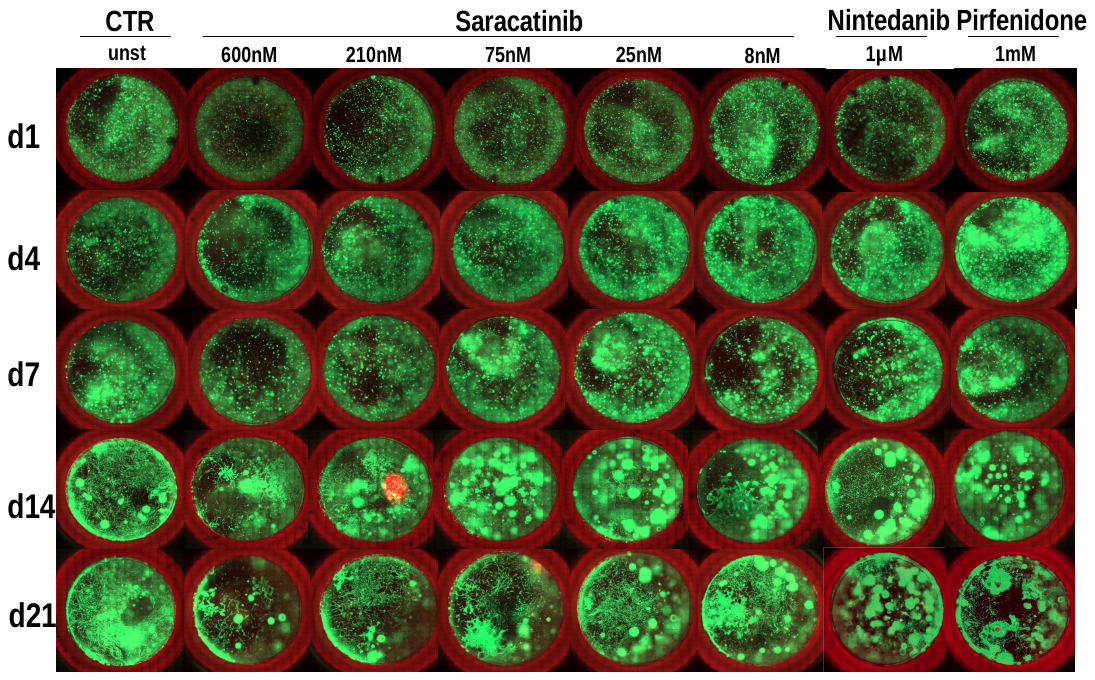

***Figure S5.*** ***Live death staining of bronchosphere cultures with IPF lung fibroblasts.*** *Green staining detects vivid cells while red staining detects dead cells. In all bronchosphere cultures w/wo treatment with saracatinib, pirfenidone or nintedanib hardly any dead cells were detected at day 4 (d4), day 7 (d7), day 14 (d14), and day 21 (d21).*

### Supplementary References

1.	American Thoracic Society. Idiopathic pulmonary fibrosis: diagnosis and treatment. International consensus statement. American Thoracic Society (ATS), and the European Respiratory Society (ERS). *Am J Respir Crit Care Med* **161**, 646-664 (2000).

2.	American Thoracic Society/European Respiratory Society International Multidisciplinary Consensus Classification of the Idiopathic Interstitial Pneumonias. This joint statement of the American Thoracic Society (ATS), and the European Respiratory Society (ERS) was adopted by the ATS board of directors, June 2001 and by the ERS Executive Committee, June 2001. *Am J Respir Crit Care Med* **165**, 277-304 (2002).

3.	Raghu, G. *et al.* An official ATS/ERS/JRS/ALAT statement: idiopathic pulmonary fibrosis: evidence-based guidelines for diagnosis and management. *Am J Respir Crit Care Med* **183**, 788-824 (2011).

4.	Shinkai, Y. *et al.* RAG-2-deficient mice lack mature lymphocytes owing to inability to initiate V(D)J rearrangement. *Cell* **68**, 855-867 (1992).

5.	Pearson, T. *et al.* Non-obese diabetic-recombination activating gene-1 (NOD-Rag1 null) interleukin (IL)-2 receptor common gamma chain (IL2r gamma null) null mice: a radioresistant model for human lymphohaematopoietic engraftment. *Clinical and experimental immunology* **154**, 270-284 (2008).

6.	Ivics, Z., Hackett, P.B., Plasterk, R.H. & Izsvak, Z. Molecular reconstruction of Sleeping Beauty, a Tc1-like transposon from fish, and its transposition in human cells. *Cell* **91**, 501-510 (1997).

7.	Garrels, W. *et al.* Germline transgenic pigs by Sleeping Beauty transposition in porcine zygotes and targeted integration in the pig genome. *PloS one* **6**, e23573 (2011).

8.	Nagai, T. *et al.* A variant of yellow fluorescent protein with fast and efficient maturation for cell-biological applications. *Nature biotechnology* **20**, 87-90 (2002).

9.	Hackett, N.R. *et al.* RNA-Seq quantification of the human small airway epithelium transcriptome. *BMC Genomics* **13**, 82 (2012).

10.	Prasse, A. *et al.* A vicious circle of alveolar macrophages and fibroblasts perpetuates pulmonary fibrosis via CCL18. *Am J Respir Crit Care Med* **173**, 781-792 (2006).

11.	Cheung, K.J., Gabrielson, E., Werb, Z. & Ewald, A.J. Collective invasion in breast cancer requires a conserved basal epithelial program. *Cell* **155**, 1639-1651 (2013).

12.	Okkelman, I.A., Dmitriev, R.I., Foley, T. & Papkovsky, D.B. Use of Fluorescence Lifetime Imaging Microscopy (FLIM) as a Timer of Cell Cycle S Phase. *PloS one* **11**, e0167385 (2016).

13.	Leonard, A.K. *et al.* Methods for the visualization and analysis of extracellular matrix protein structure and degradation. *Methods Cell Biol* **143**, 79-95 (2018).

14.	Kurita, Y. *et al.* Pirfenidone inhibits myofibroblast differentiation and lung fibrosis development during insufficient mitophagy. *Respiratory research* **18**, 114 (2017).

15.	Martin, M. Cutadapt removes adapter sequences from high-throughput sequencing reads. *2011* **17**, 3 (2011).

16.	Parekh, S., Ziegenhain, C., Vieth, B., Enard, W. & Hellmann, I. zUMIs - A fast and flexible pipeline to process RNA sequencing data with UMIs. *Gigascience* **7** (2018).

17.	Zerbino, D.R. *et al.* Ensembl 2018. *Nucleic Acids Res* **46**, D754-D761 (2018).

18.	Dobin, A. *et al.* STAR: ultrafast universal RNA-seq aligner. *Bioinformatics* **29**, 15-21 (2013).

19.	Durinck, S., Spellman, P.T., Birney, E. & Huber, W. Mapping identifiers for the integration of genomic datasets with the R/Bioconductor package biomaRt. *Nat Protoc* **4**, 1184-1191 (2009).

20.	Waltman, L. & van Eck, N.J. A smart local moving algorithm for large-scale modularity-based community detection. *The European Physical Journal B* **86**, 471 (2013).

21.	Butler, A., Hoffman, P., Smibert, P., Papalexi, E. & Satija, R. Integrating single-cell transcriptomic data across different conditions, technologies, and species. *Nature biotechnology* **36**, 411-420 (2018).

22.	Adams, T.S. *et al.* Single-cell RNA-seq reveals ectopic and aberrant lung-resident cell populations in idiopathic pulmonary fibrosis. *Sci Adv* **6**, eaba1983 (2020).

23.	McInnes, L., Healy, J. & Melville, J. in arXiv e-prints (2018).

24.	Kuleshov, M.V. *et al.* Enrichr: a comprehensive gene set enrichment analysis web server 2016 update. *Nucleic Acids Res* **44**, W90-97 (2016).

25.	Kanehisa, M. & Goto, S. KEGG: kyoto encyclopedia of genes and genomes. *Nucleic Acids Res* **28**, 27-30 (2000).

26.	Prasse, A. *et al.* BAL Cell Gene Expression Is Indicative of Outcome and Airway Basal Cell Involvement in Idiopathic Pulmonary Fibrosis. *Am J Respir Crit Care Med* **199**, 622-630 (2019).

27.	Subramanian, A. *et al.* A Next Generation Connectivity Map: L1000 Platform and the First 1,000,000 Profiles. *Cell* **171**, 1437-1452.e1417 (2017).

28.	Yu, G. *et al.* Matrix metalloproteinase-19 is a key regulator of lung fibrosis in mice and humans. *Am J Respir Crit Care Med* **186**, 752-762 (2012).

29.	Philipp, F. *et al.* Human Teratoma-Derived Hematopoiesis Is a Highly Polyclonal Process Supported by Human Umbilical Vein Endothelial Cells. *Stem Cell Reports* **11**, 1051-1060 (2018).

30.	Ashcroft, T., Simpson, J.M. & Timbrell, V. Simple method of estimating severity of pulmonary fibrosis on a numerical scale. *J Clin Pathol* **41**, 467-470 (1988).

31.	Huleihel, L. *et al.* Modified mesenchymal stem cells using miRNA transduction alter lung injury in a bleomycin model. *Am J Physiol Lung Cell Mol Physiol* **313**, L92-L103 (2017).

32.	Elmore, S.A. *et al.* in Toxicol Pathol, Vol. 44, Edn. 2016/02/18 173-188 (2016).
